# Probing molecular specificity with deep sequencing and biophysically interpretable machine learning

**DOI:** 10.1101/2021.06.30.450414

**Authors:** H. Tomas Rube, Chaitanya Rastogi, Siqian Feng, Judith F. Kribelbauer, Allyson Li, Basheer Becerra, Lucas A. N. Melo, Bach Viet Do, Xiaoting Li, Hammaad H. Adam, Neel H. Shah, Richard S. Mann, Harmen J. Bussemaker

## Abstract

Quantifying sequence-specific protein-ligand interactions is critical for understanding and exploiting numerous cellular processes, including gene regulation and signal transduction. Next-generation sequencing (NGS) based assays are increasingly being used to profile these interactions with high-throughput. However, these assays do not provide the biophysical parameters that have long been used to uncover the quantitative rules underlying sequence recognition. We developed a highly flexible machine learning framework, called ProBound, to define sequence recognition in terms of biophysical parameters based on NGS data. ProBound quantifies transcription factor (TF) behavior with models that accurately predict binding affinity over a range exceeding that of previous resources, captures the impact of DNA modifications and conformational flexibility of multi-TF complexes, and infers specificity directly from *in vivo* data such as ChIP-seq without peak calling. When coupled with a new assay called Kd-seq, it determines the absolute affinity of protein-ligand interactions. It can also profile the kinetics of kinase-substrate interactions. By constructing a biophysically robust foundation for profiling sequence recognition, ProBound opens up new avenues for decoding biological networks and rationally engineering protein-ligand interactions.

## Introduction

Gene regulatory and signal transduction networks rely on sequence-specific molecular recognition to guide constituent proteins to preferentially bind to or chemically modify specific nucleic-acid or amino-acid ligands or substrates. These interactions often span orders of magnitude in strength and are modulated not only by sequence, but also by other *in vivo* effects such as competition, cooperation, saturation and chemical modifications^1^. As even weak ligands can be functional^2–4^, comprehensive and accurate profiling of sequence recognition is essential to decode these networks.

Sequence-specific interactions are most appropriately described in terms of biophysical parameters such as equilibrium constants and reaction rates. Sequence recognition models, which often take the form of position-specific scoring matrices^5^, encode how a protein recognizes any sequence and have proven useful for predicting binding targets and the impact of genetic variation^1^. However, in their current form, they fall short of predicting actual biophysical constants. To build truly quantitative recognition models, we need improved algorithms along with high-quality datasets to train them.

In recent years, NGS has dramatically increased the throughput with which molecular interactions can be probed. In particular, high-throughput methods coupling NGS with in *vitro* selection on random ligand pools have emerged as powerful and flexible tools for the unbiased profiling of sequence recognition. This includes SELEX methods for TFs^6–16^ and RNA-binding proteins^17,18^, as well as protein display methods for proteases^19^ and T-cell receptors^20^. Transforming the resulting sequencing reads into quantitative recognition models remains challenging, as the biophysical properties are only indirectly encoded in the sequencing reads. Moreover, randomized ligand pools can be extremely complex and even the best sequences can go unobserved. There is currently no general method that systematically addresses these issues.

Here, we solve this problem with a flexible machine learning framework, called ProBound, capable of learning biophysically interpretable recognition models from a wide range of sparse NGS data. It can quantify relative affinities, absolute dissociation constants, cooperativity, methylation sensitivity, and enzymatic parameters by analyzing data from various *in vivo* or *in vitro* assays covering DNA, RNA, or protein ligands. The resulting binding models are highly accurate, as illustrated by their superior performance relative to existing resources. While current methods support elements of these features^21–25^, ProBound allows for unprecedented quantitative rigor and generality.

## Results

### The ProBound framework

ProBound uses three layers to systematically model NGS data (Figure 1; Methods): a *binding layer* that predicts the binding free energy or enzymatic efficiency from sequence; an *assay layer* that predicts the post-selection frequency of a ligand; and a *sequencing layer* that represents the stochastic sampling of DNA reads during deep sequencing. Together, these elements are combined in a likelihood function that aims to explain the observed distribution of read counts across multiple selection rounds or conditions in terms of the sequence features of the ligand. Each layer is easily extensible; for example, the binding layer can model TF complexes by accommodating multiple recognition models and their interactions. Flexibility in the assay layer enables the modeling of alternative selection processes (e.g. catalysis) and the utilization of multiple assays to measure more complex phenomena (e.g. cooperativity).

**Figure 1:**
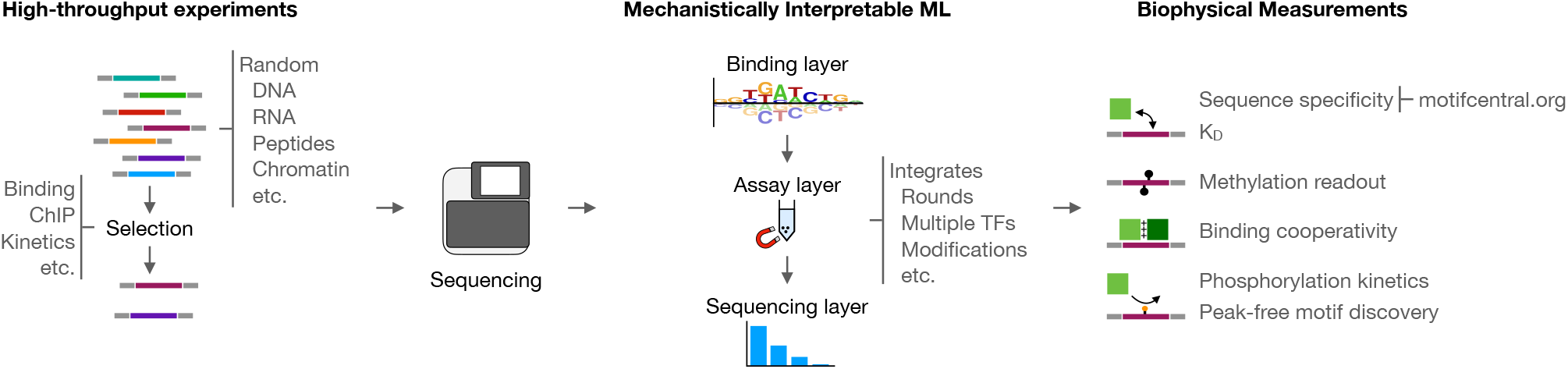
Overview of the ProBound method. A range of high-throughput experiments utilize selection on random DNA, RNA and displayed protein libraries coupled with NGS to characterize sequence-specific molecular interactions. ProBound uses machine learning tailored to model the recognition, selection, and sequencing processes in such experiments to infer biophysically meaningful sequence recognition models from a wide range of NGS data.

### A compendium of accurate TF binding models

Our initial objective was to analyze thousands of published SELEX datasets^9,10,12,14,15,26–28^ and produce high-quality TF binding models that capture low-affinity binding, an important yet difficult-to-detect gene regulatory phenomenon^2–4, 22^. This required us to quantify TF sequence recognition over a wide affinity range, rather than merely classify sequences as “bound” or “unbound”. We therefore assembled a training database of 2,124 published SELEX datasets and designed a computational pipeline to uniformly build binding models (Figure 2a; Supplemental Table 1; Methods). To assess the generalization performance of our models, we linked each TF to published protein binding microarray (PBM), ChIP-seq, and non-training SELEX data. We computed three complementary performance metrics: meaningful affinity fold-range (MAFR), a new metric that provides a conservative bound on the ability of a model to detect low-affinity binding; *R*^2^, the fraction of signal variance explained by the model; and area under the precision-recall curve (AUPRC), a common metric^22,25,29,30^ for quantifying how well a model classifies genomic regions as bound or unbound as determined by ChlP-seq peaks^31^. We used these to benchmark our models to those in major resources, linking all JASPAR^32^, DeepBind^30^, HOCOMOCO^33^, and Jolma et al. (2013^)26^ models by TF. On average, ProBound outperformed these resources across all metrics (Figure 2b), with the PBM and SELEX metrics displaying the largest improvement. The less notable improvement in AUPRC is likely due to bias towards high-affinity sequences in ChlP-seq peaks, for which accurate low-affinity predictions are less relevant^22^. Below, we will introduce an alternative method for analyzing ChIP-seq data that eliminates the need for ChIP-seq peak discovery.

**Figure 2:**
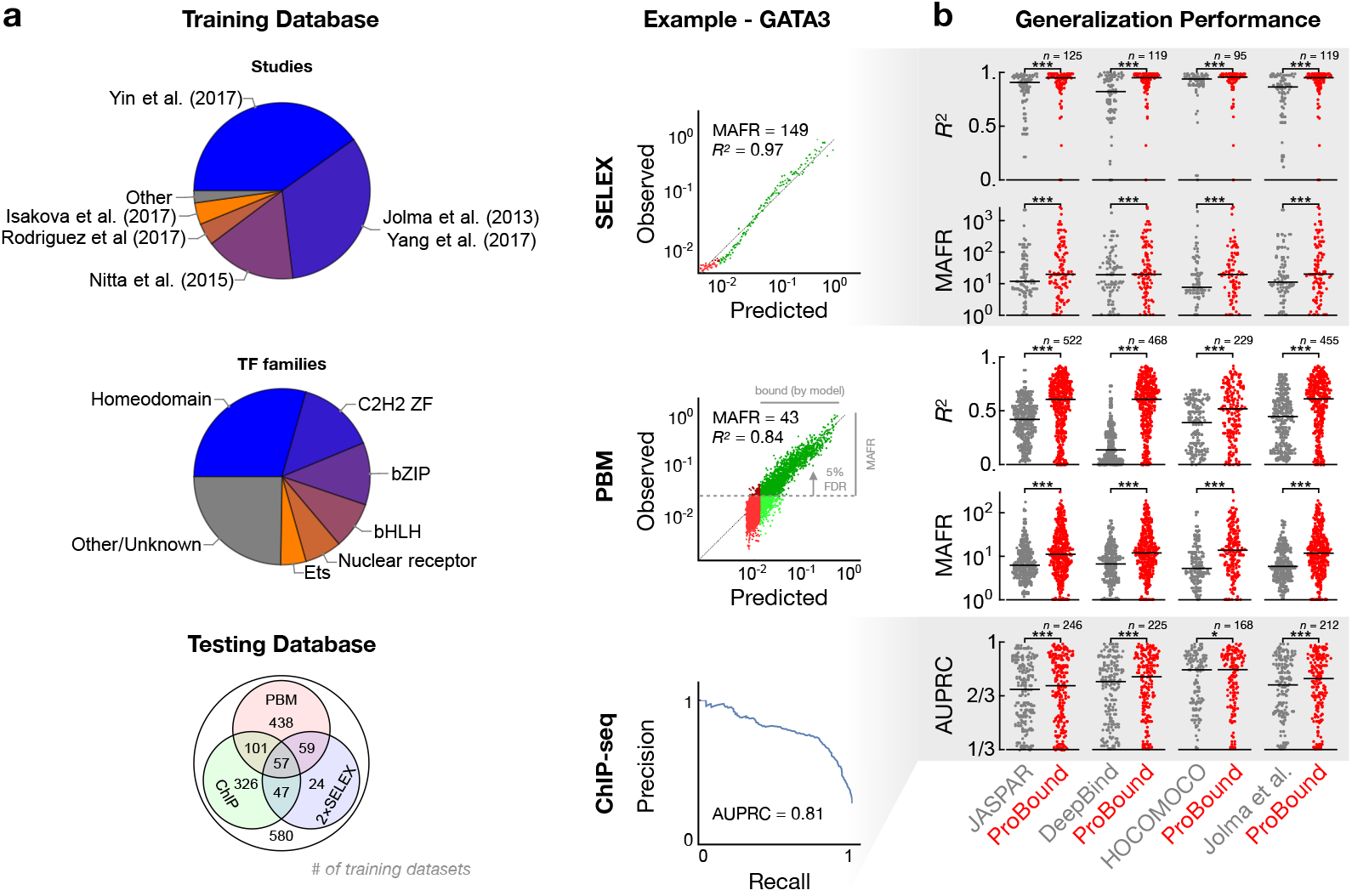
Validation of TF binding model performance. **(a)** Breakdown of the training dataset used to build recognition models by originating study and TF family (pie charts) and by availability of testing data used to evaluate them (Venn diagram). Representative SELEX (top) and PBM (middle) comparisons of observed and model-predicted binding signals used to quantify generalization performance. Each point in the scatter plots corresponds to either 500 SELEX probes or 10 PBM probes; green indicates where the model predicts binding above an estimated baseline (see Methods), while darker points indicate the meaningful affinity fold-range (MAFR) of observed binding signal over which at most 5% of predicted binding was below the baseline. Representative precision-recall curve (bottom) for the ChIP-seq peak classification task used to quantify model performance in terms of AUPRC. **(b)** Performance comparison of ProBound models vs. popular existing resources. For each ProBound and resource model pair (points), the average score was computed for all matching testing datasets. Horizontal bars indicate median performance. Significance was computed using the Wilcoxon signed-rank test.

Over the years, a number of TFs have been assayed many times by different research groups and SELEX platforms. We reasoned that jointly analyzing such data would produce a “consensus” model focused on the true binding signal rather than platform-specific biases (Figure S1a). Encouragingly, such consensus models displayed significantly improved performance when compared to traditional single-experiment models (Figure S1b), indicating that multi-experiment analysis can improve binding predictions. Finally, to facilitate adoption by other researchers, we have made a curated version of our models, comparative analyses, and computational tools readily available through a comprehensive resource at motifcentral.org.

### Quantifying TF binding cooperativity

Variables beyond sequence, such as co-factor interactions and DNA methylation, significantly influence TF behavior *in vivo*, and therefore, TF binding models must account for them in order to improve binding predictions. We first focused on co-factors, which modulate TF binding in a cell-type-specific manner. Despite the growing number of SELEX assays characterizing TF complexes^9,11,34^, it remains a challenge to quantify sequence recognition in a way that clearly separates the contributions from many potential TF complexes and their various internal structural configurations – a problem that grows exponentially with the number of factors assayed. In a novel approach that builds upon our multi-experiment framework, we measure subunit binding specificity and cooperativity by explicitly modeling the allowed complexes in multiple SELEX datasets that probe different TF combinations.

We first applied this method on the complex formed by three highly conserved *Drosophila* homeodomain proteins: Homothorax (Hth), Extradenticle (Exd) and Ultrabithorax (Ubx). Previous studies showed that Ubx and Exd form fixed-spacer heterodimers^10,22^ and that Hth uses multiple relative spacings to bind cooperatively with similar heterodimers^34^. To characterize Hth:Exd:Ubx, we first performed SELEX-seq with all three factors and then analyzed these data in conjunction with our previous monomer and heterodimer data (Figure 3a, S2a). Importantly, we modeled the ternary complex with two subunits representing Hth and Exd:Ubx; the total binding energy was the sum of their independent binding specificities and of a cooperativity term that depended on their relative position and orientation.

**Figure 3:**
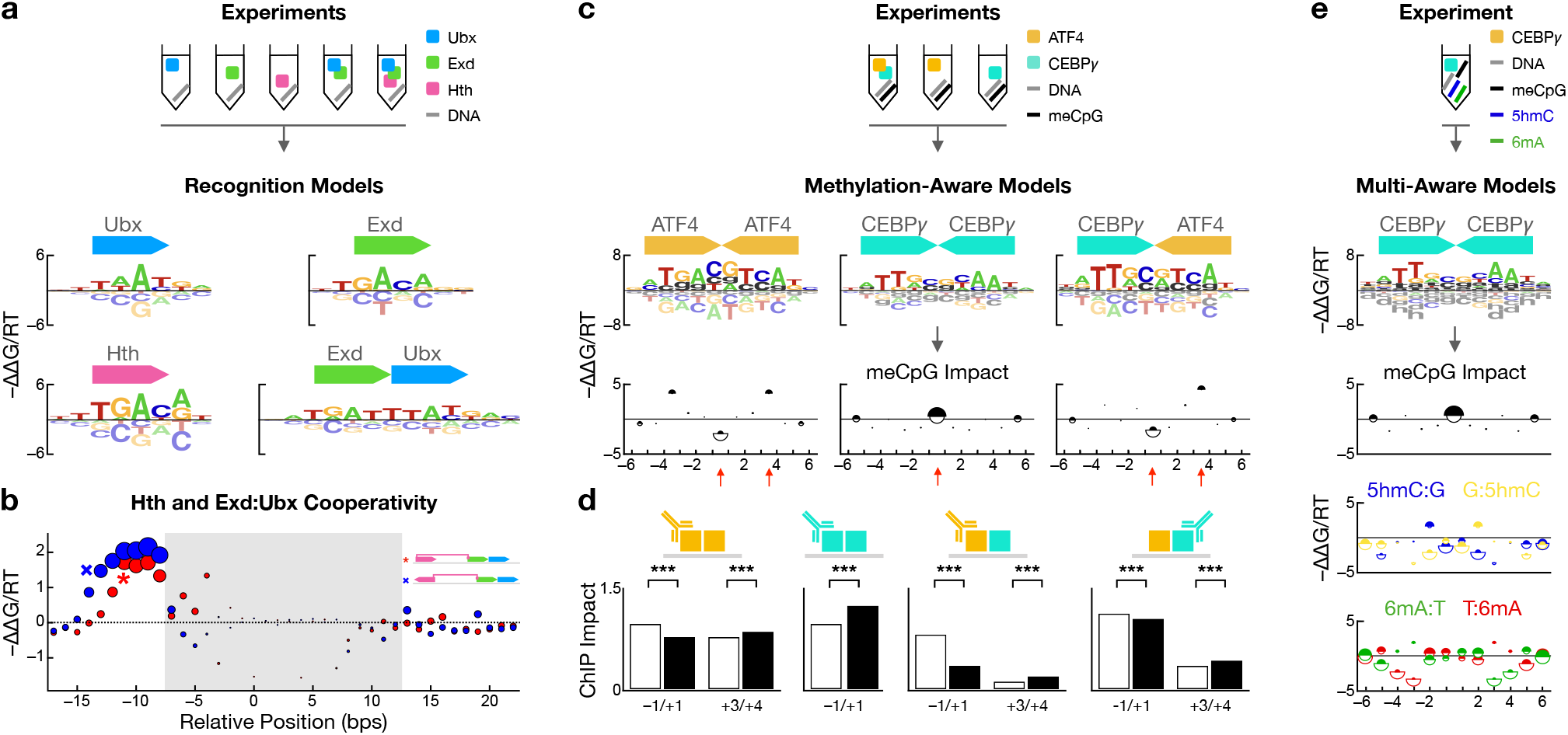
Integrated modeling of complementary assays quantifies the impact of methylation and co-factors on TF binding. **(a)** Combinations of TFs assayed (top) and unified model learned by ProBound (bottom). The model consists of the inferred energy logos for the monomeric and dimeric complexes (motifs) and the **(b)** inferred binding cooperativity (y-axis) between Hth and Exd:Ubx for different relative positions (x-axis) and orientations (red: parallel; blue: anti-parallel) of the subunits. Disψreas proportional to the affinity of the strongest predicted sequence highlight the most stable configurations. Shaded region indicates overlapping motifs. Schematics (inset) illustrate two configurations indicated on the plot. **(c)** Combinations of TFs and methylated/unmethylated libraries assayed (schematic); methylation-aware binding models (motifs) using the alphabet in Figure S3a; and impact of meCpG on binding free energy (plots; −ΔΔ*G*_CpG→meCpG_/*RT* on y-axis) as a function of position within the binding site (x-axis). Half-disk areas are proportional to the maximum affinity when either CpG (white) or meCpG (black) is substituted at the corresponding position in the highest-affinity sequence and highlight positions with high-affinity methylation readout. **(d)** Impact of substituting a CpG (white) or meCpG (black) at a specific position in the highest-affinity binding site as quantified using ChIP-seq data. Each pair of bars corresponds to a substitution at a specific position and to red arrows in (c). Antibody symbols indicate respective immunoprecipitated factor. Asterisks indicate significance computed using an *F*-test (see Methods and Supplemental Table 2). **(e)** Same as (c) for data simultaneously measuring methylation readout for meCpG, 5hmC, and 6mA modifications.

The resulting model revealed significant cooperativity (ΔΔ*G*_config_ ≈ 2*RT*) when Hth binds 8-13 bps upstream of Exd:Ubx (Figure 3b), which, along with our monomer and heterodimer models, mirrored our previous results^22, 34^. While a larger spacing is tolerated when Hth is reversed, cooperativity is lost when Hth binds far away from the Exd:Ubx half-site, regardless of orientation. As expected, selection in the Hth-Exd-Ubx experiment was driven by multiple subcomplexes with alternate spacing preferences (Figure S2b), underscoring the need to simultaneously model all preferences. As a further test, we reanalyzed published data for POU2F:GSC2 and GCM1:ELK1 in combination with matched monomer data^11,26^. In both cases, strong binding cooperativity was detected at a specific relative offset (Figure S2c, d).

### Learning methylation-aware TF binding models

Next, we focused on another variable affecting *in vivo* binding: DNA methylation. Chemical modifications to DNA, such as fully methylated CpG dinucleotides (meCpG), are common epigenetic marks that can alter TF binding, and thus, gene regulation^35–38^. Unlike existing methods that compare methylated and normal SELEX libraries to detect TF “methylation readout” at the level of enriched subsequences^14,16,39^, we used ProBound with an extended alphabet (Figure S3a, Methods) and our multi-experiment framework to learn methylation-aware binding models that resolve the position-specific impact of methylation (ΔΔ*G*_CpG→meCpG_), enabling binding predictions to any (un)methylated sequence.

We tested this approach by simultaneously uncovering the effect of meCpG on the ATF4:CEBP*γ* heterodimer while controlling for the confounding influence of their respective homodimers. Using data for all combinations of ATF4/CEBP*γ* and normal/methylated DNA (Figure S3b), we simultaneously learned methylation-aware binding models for all three dimers (Figure 3c, Methods). These predict methylation induced stabilization/destabilization patterns (Figure 3c, S3c) consistent with previous analyses of the ATF4 homodimer^15^ and similar to those of the related CEBP*β* homodimer^15^ and ATF4:CEBP*β* heterodimer^39^. Strikingly, ATF4 overrides CEBPχ to retain its methylation readout at the central position of the heterodimer complex. Importantly, we used ChIP-seq data to estimate the impact of these position-specific methylation sensitivities *in vivo*, and found that methylation significantly affected binding in the direction predicted by our models (Figure 3d, Methods).

Other DNA modifications, such as N^6^-methyladenine (6mA) and 5-hydroxymethylcytosine (5hmC), can also be functional^40–45^. To characterize their impact, we extended the EpiSELEX-seq protocol to assay multiple sub-libraries simultaneously: unmethylated, meCpG, 5hmC, and 6mA (Figure 3e and S4a). Not only is this simpler than assaying each methylation mark separately, it also reduces experimental error. Repeating the binding assay for CEBP*γ* and jointly analyzing all four libraries reveals significant and distinct stabilization/destabilization patterns for both 5hmC and 6mA (Figure 3e and S4b). Notably, the inferred meCpG methylation sensitivity is identical to what we found above. These results illustrate both the scalability of our approach and the impact 5hmC and 6mA can have on binding.

### Measuring absolute binding constants using SELEX

While we have focused on quantifying binding specificity in terms of relative affinities, knowledge of *absolute* affinities is necessary for predicting equilibrium occupancy and for comparing different TFs on a common scale. Fundamentally, SELEX assays probe *relative* ligand frequencies, and so far, have only been used to estimate *relative* affinities. To overcome this limitation, we developed a novel assay called *K_D_*-seq. It uses ProBound to jointly analyze the input, bound, and free probes from a selection round and produce both a specificity model and an estimate of the absolute dissociation constant (*K_D_*) for a reference sequence. Intuitively, *K_D_*-seq uses a sum rule to relate the relative ligand frequencies of the three libraries and convert them to absolute binding probabilities (Figure 4a, Methods).

**Figure 4:**
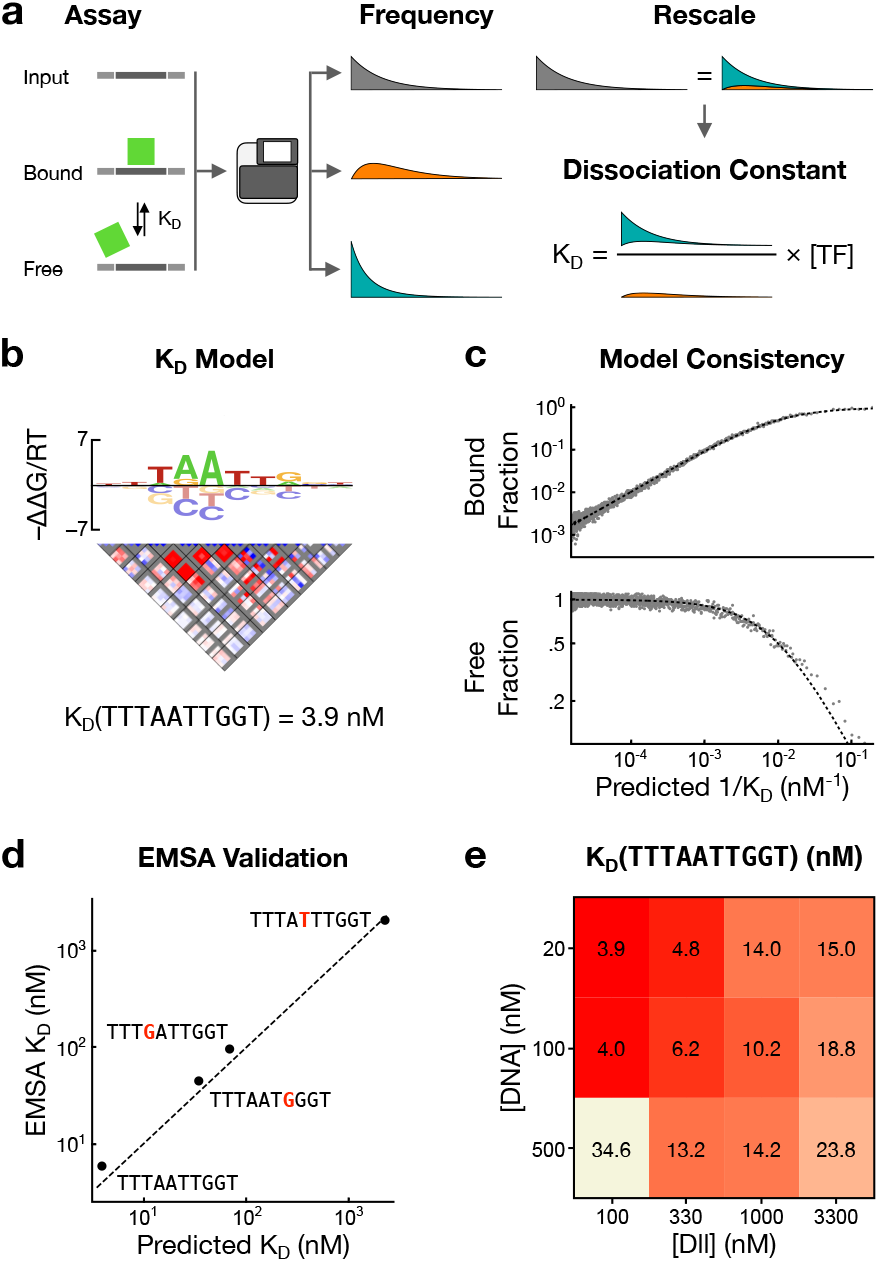
ProBound infers absolute *K_D_* values. **(a)** Schematic overview of the *K_D_*-seq method. After a TF is incubated with a randomized DNA library, the bound, free, and input probes are sequenced, measuring the relative probe frequencies in each fraction. This can be used to estimate the absolute binding probabilities (and hence *K_D_*) with a sum rule that relates the three frequencies. **(b)** *K_D_* model for Dll consisting of a specificity model with an energy logo (top) and an interaction matrix (middle), which together predict the relative binding affinity, and the absolute *K_D_* for a reference sequence (bottom). The interaction plot shows stabilizing (red) and destabilizing (blue) corrections to the energy logo for each pair of positions (boxes) and bases (pixels) lin the logo. Gray indicates prohibited corrections. Model generated from data where [Dll] = 100nM and [DNA] = 20nM. **(c)** Comparison of the predicted 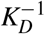 (x-axis) and observed probe fractions (y-axis) in the bound (top) and free (bottom) libraries. Points represent the average observed fraction for 500 probes binned by predicted *K_D_*. Dashed line indicates expected value assuming equilibrium binding model. **(d)** Comparison between EMSA-measured (y-axis) and model-predicted (x-axis) *K_D_* values for four probes. **(e)** *K_D_* of the sequence TTTAATTGGT as estimated by *K_D_*-seq for different Dll and DNA concentrations.

We initially tested *K_D_*-seq using the *Drosophila* homeodomain protein Distal-less (Dll) at low DNA and TF concentrations (100nM and 20nM, respectively) to achieve strong enrichment and avoid excessive binding saturation.

The resulting model (Figure 4b) accurately predicted enrichment in the bound and free libraries over three orders of magnitude in *K_D_* (Figure 4c). For validation, we measured the *K_D_* values of the optimal model-predicted binding site and three suboptimal sequences using EMSA and found excellent quantitative agreement (Figure 4d). We then confirmed the robustness of *K_D_*-seq affinity measurements by repeating the assay at different TF and DNA concentrations (Figure S5a). The resulting specificity models were virtually identical (pairwise *r*^2^ for ΔΔ*G* ranging from 0.974-0.998), with the fraction of bound DNA changing as expected (Figure S5b). While the estimated *K_D_* of the highest-affinity sequence was robust in many conditions, it shifted at extremely high TF concentrations (~600-fold above EMSA-measured *K_D_*) or when DNA concentration was significantly above that of the TF (Figure 4e).

ProBound can also learn *K_D_* models by jointly analyzing the bound and input libraries of multiple SELEX experiments at different TF concentrations. Intuitively, this approach leverages saturation effects to determine the absolute affinity scale. For Dll, the *K_D_* models from the two approaches are very similar (Figure S5a,c-d). When applied to multi-concentration RNA Bind-N-seq^18^ data for RBFOX2, the resulting *K_D_*-model captured the observed transition from linear to saturated selection in the experiments (Figure S5f). Finally, we note that ProBound can estimate relative affinities using only the free and bound libraries, as in the Spec-seq^46^ assay (Figure S5e).

### Peak-free motif discovery from ChIP-seq data

While the preceding analyses have focused on quantifying the impact of co-factors and TF concentration on *in vitro* binding, we also wished to learn their *in vivo* impact directly from ChIP-seq data. Standard motif discovery algorithms aim to discover overrepresented sequences within discrete genomic regions – identified by “peak callers” – that harbor a statistically significant enrichment of ChIP-seq reads. Peak calling is useful for identifying the most prominent genomic binding sites, but it ignores information about cis-regulatory logic contained within more weakly bound regions. We hypothesized that by directly modeling the enrichment between the input and ChIP libraries, ProBound can extract such information even from weakly enriched regions.

To test this approach, we used ProBound to discover the factors driving the selection in glucocorticoid receptor (GR) ChIP-seq data from the IMR90 cell line^47^ (see Methods). It found four binding models: one consistent with the GR consensus sequence^49,50^ and three others consistent with known GR co-factors AP-1, FOXA1, and TEAD^47,51^ (Figure 5a). Inspired by our multi-concentration analysis above, we next set out to quantify the impact the nuclear concentration of a TF can have on binding. We did so by jointly analyzing multiple ChIP-seq datasets that probe GR binding in the murine hippocampus after treatment with varying levels of corticosterone (CORT)^48^, an agonist that increases the nuclear concentration of GR (Figure 5b). The resulting model captured sample-specific activity parameters reflective of GR nuclear concentration that were proportional to CORT concentration (Figure 5b).

**Figure 5:**
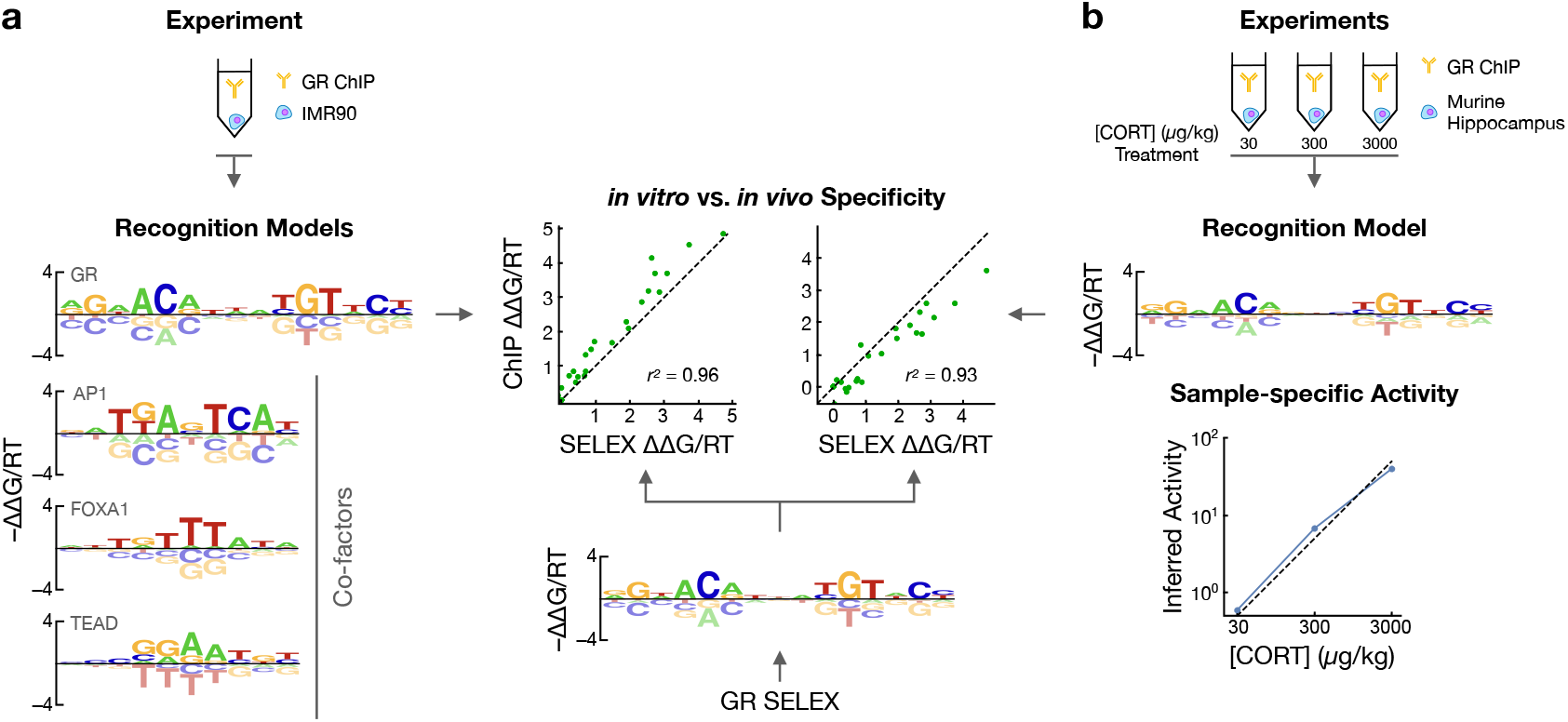
ProBound learns quantitative binding models and sample-specific activities using peak-free ChIP-seq analysis. **(a)** Binding models for GR and three co-factors (left) learned from GR ChIP-seq data from the IMR90 cell line^47^ and for GR from a SELEX dataset (center). The scatterplot compares the energy coefficients learned from ChIP-seq (y-axis) and SELEX (x-axis) data^9^. **(b)** Combined specificity (top) and sample-specific binding activity (bottom) model learned by jointly analyzing three GR ChIP-seq datasets after treatment with 30, 300, or 3000*μ*g/kg of corticosterone (CORT)^48^. The scatterplot (left) compares the energy coefficients as in (a).

It should be noted that both these models were constructed on data that was intentionally downsampled to less than one mapped read per kb of genomic sequence on average. Thus, even when peak discovery is ineffective, ChIP-seq data clearly contain sufficient information to reliably infer TF binding models, capture the influence of co-factors, and quantify biologically meaningful cell state parameters. Significantly, the free-energy parameters of both GR binding models showed striking agreement with those from a model trained on *in vitro* data^9^ (*r*^2^ = 0.96 and *r*^2^ = 0.93, respectively; Figure 5a, b), suggesting that *in vitro* and *in vivo* observations of binding specificity can, in fact, be highly concordant.

### Profiling tyrosine kinase kinetics

Biological processes that employ sequence-specific protein-protein interactions are increasingly being studied with display assays utilizing diverse DNA-templated protein libraries^19,20,52^. While these methods are profiling these interactions more comprehensively than ever before, interpreting the data remains challenging for many of the same reasons as above. Furthermore, current analytical methods tend to focus on detecting enriched sequence features rather than explicitly estimating binding constants or enzymatic parameters. Given the similarities with SELEX assays, we were motivated to use ProBound to characterize protein sequence recognition.

As a proof-of-concept, we focused on a process critical to many signal transduction pathways in the cell – the phosphorylation of tyrosine residues on proteins. Recently, the substrate sequence preferences of several tyrosine kinases were surveyed with a bacterial display library containing thousands of known kinase targets^53^. To comprehensively profile the preferences for one of these kinases, c-Src, in an unbiased way, we repeated the assay with a new library design that randomizes ten amino-acid residues around a fixed central tyrosine and exposed this library to c-Src for varying durations (Figure 6a; Methods). After sequencing, we jointly analyzed all time points to learn a model that predicts the sequence-specific catalytic efficiency *k*_eff_, a simple metric that is often used to compare different substrates against a single enzyme. Visualizing the inferred efficiency model as a sequence logo (Figure 6b) revealed a position-specific pattern of favorable residues that were consistent with the earlier study^53^.

**Figure 6:**
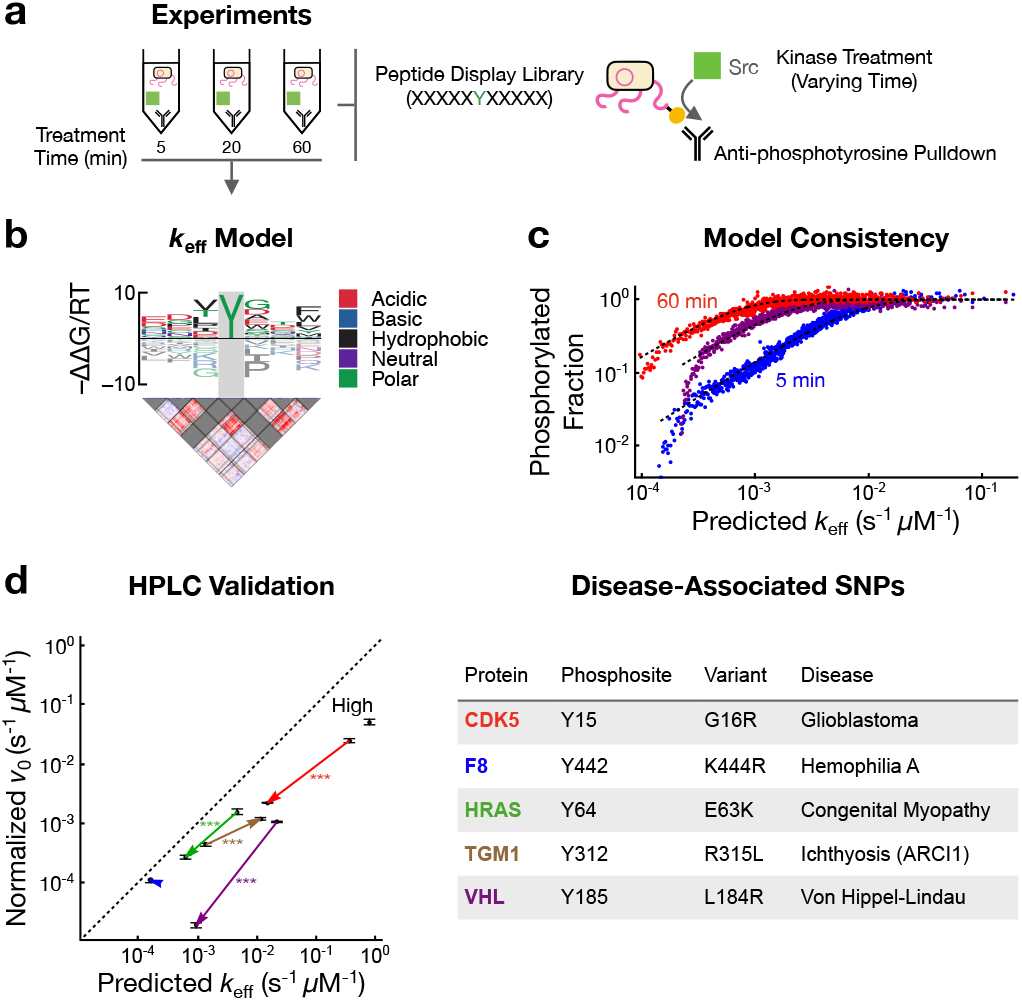
ProBound quantifies sequence-dependent kinetics of the tyrosine kinase c-Src. **(a)** Schematic overview of the bacterial display assay used profile the sequence specificity of the tyrosine kinase c-Src. **(b)** *k*_eff_ model for c-Src with an energy logo (top) and an interaction matrix (bottom) trained on data from 5, 20 and 60 minutes of exposure. The central position of the model was fixed to recognize tyrosine (gray). **(c)** Comparison of the predicted *k*_eff_ (x-axis) and phosphorylated fraction (y-axis) for 5 (blue), 20 (purple) and 60 minutes (red) of exposure to c-Src. Points represent the average observed phosphorylated fraction for 500 probes binned by predicted *k*_eff_. Dashed lines indicate expected value according to model. **(d)** Comparison of the HPLC-measured normalized initial phosphorylation rate *v*_0_ (y-axis, *n* = 3 technical replicates) and the model-predicted *k*_eff_ (x-axis) for five disease-associated WT/MUT SNP pairs (arrows) and a peptide predicted to have high activity (table and Supplemental Table 3). The concentration of c-Src was 500 nM and that of the substrate peptide was 100 *μ*M. Error bars indicate the standard error of the mean and asterisks indicate significance computed using a two-sided *t*-test.

The model also accurately captures the observed fraction of phosphorylated peptides over a 100-fold range in *k*_eff_ for all three time points (Figure 6c).

To validate the model, we used high-performance liquid chromatography (HPLC) to measure the phosphorylation rates for eleven peptides. As genetic variants can significantly impact phosphorylation rates^54^, we used the PTMVars database^55^ to find four disease-associated SNPs predicted by our ProBound model to have a large allelic difference. Indeed, measurements of their normalized initial phosphorylation rate differed significantly in the direction predicted by the model (Figure 6d). In addition, there was no measurable difference for a SNP predicted to cause only a small allelic difference for the F8 protein, and a model-generated high-efficiency peptide (Src-high) was indeed the highest. Significantly, these predictions tracked measurements over three orders of magnitude in *k*_eff_, suggesting that ProBound is a powerful new tool for quantifying enzyme specificity.

## Discussion

A major goal of this study was to rigorously estimate biophysical parameters from NGS data using machine learning. While biochemists have measured such parameters for decades, these measurements are generally low-throughput. By contrast, high-throughput sequencing-based analysis tends to focus on the detection of enrichment patterns that only indirectly reflect these quantities. Moreover, modern machine learning methods such as deep neural networks tend to yield highly overparametrized black box models whose parameters have no direct biophysical meaning. Here, we showed that by explicitly modeling the assay process, we can use machine learning to turn DNA sequencers into virtual measurement devices that accurately quantify biophysical parameters. Molecular biologists and computer scientists often address the same question using very different language; for instance, classifier performance and binding free energies are both used to quantify sequence recognition. We hope that approaches such as ours help keep the literature more coherent and inspire direct experimental validation of algorithm performance.

Central to our approach is the observation that some quantities cannot be estimated through pairwise enrichment analysis but only through more structured integration of complementary data. One example is our combinatorial approach to the separation of different TF complexes, which we also extended to methylation-aware binding models. Another is how analyzing the bound, free, and input fractions jointly – not pairwise – allows absolute affinities to be measured. Our approach is reminiscent of more traditional biochemical assays, which collect data across different time points, concentrations, or fractions, and use curve fitting to estimate constants. As we study increasingly complex aspects of sequence recognition — such as the combined impact of sequence, co-factors, DNA methylation, and TF concentrations, or the integration of *in vitro* and *in vivo* perspectives — we foresee that rigorous integration of complementary data along the lines that we have sketched here will become increasingly important. More generally, we anticipate that the accurate and unbiased profiling of sequence recognition that ProBound enables will have numerous applications in areas of biotechnology where the rational engineering of ligands is critical.

## Methods

### Overview of the algorithm

For each experiment, the data consists of a count table enumerating the probes in each SELEX round. The core of the algorithm is a statistical model of the experiment that defines the likelihood of a set of model parameters given the count table. On a high level, this likelihood is computed by first defining the probability that each probe is bound in terms of its sequence, then predicting the probe frequencies in each library using a cumulative selection function, and finally modeling the stochastic sampling of sequencing. The model parameters are estimated from the data through numerical maximization of the likelihood.

#### Probabilistic motivation of the binding model

The binding model defines the probability that a probe is bound:

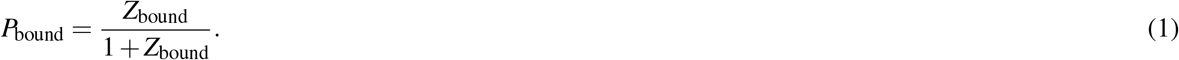

Here *Z*_bound_ is the partition function, which can be thought of as a weighted sum over microscopic states. Assuming that at most two protein molecules are bound to the probe, the partition function is given by

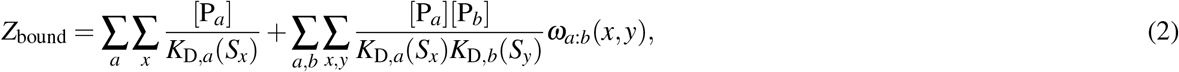

where *a* is an index that denotes protein type, [P_*a*_] is the concentration of protein *a, S_x_* a probe subsequence of length *L_a_* starting at an offset and strand denoted by *x, K*_D,*a*_(*S_x_*) is the dissociation constant for protein *a* binding *S_x_*, and ***ω***_*a:b*_(*x*_1_, *x*_2_) quantifies the cooperativity between factors *a* and *b* binding at position *x*_1_ and *x*_2_, respectively. Note that ***ω***_*a:b*_(*x*_1_, *x*_2_) equals 1 if *a* and *b* bind independently from each other, equals 0 for prohibited conformations, and is greater than 1 if the factors bind cooperatively.

It is convenient to express *K_D_* in terms of its value for a references sequence *S*_0_ and a modifier factor called the relative binding affinity:

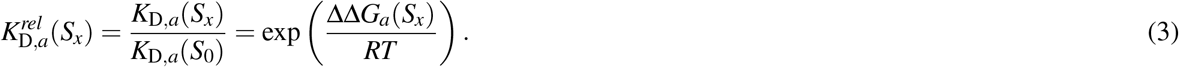

Here ΔΔ*G_m_*(*S*) ≡ Δ*G*(*S*) – Δ*G*(*S*_0_) is the difference in free-energy penalty Δ*G* of binding between *S* and *S*_0_, *R* denotes the ideal gas constant and *T* is the absolute temperature.

A central goal of our algorithm is to learn how ΔΔ*G_m_*(*S*) depends on the sequence. ProBound models this as a sum of additive contributions associated with sequence features *ϕ*:

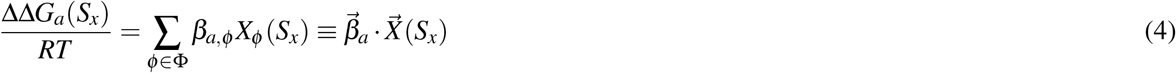

Here Φ is the set of sequence features, *β_ϕ_* is the energetic impact of *ϕ*, and *X_ϕ_*(*S_x_*) is a binary indicator of whether sequence *S_x_* contains *ϕ*. By default, Φ is simply the letter sequence along *S_x_*, meaning 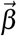 encodes aposition-specific affinity matrix (PSAM) with size matching the length of *S_x_*. ProBound can also include letter pairs (both adjacent and non-adjacent) as features.

#### Implementation of binding model

While the above derivation provides a motivation for the binding model, it has to be adapted for SELEX experiments. First, it is clear from Eq. 2 that the protein concentration [P_*a*_] and binding constant *K*_D,*a*_(*S*_0_) for a given factor *a* cannot be separately estimated from the data, but only the ratio *α_a_* = [P_*a*_]/*K*_D,*a*_(*S*_0_) can, a quantity we call the binding mode activity. We similarly define the binding mode interaction activities as *α_a:b_* = [P_*a*_] [P_*b*_]/*K*_D,*a*_(*S*_0_)*K*_D,*b*_(*S*_0_). Second, because the free protein concentration can vary between SELEX rounds *r*, the activities can take independent values in each round. Third, most experiments are performed in a low-protein-concentration regime where *Z*_bound_ ≪ 1 and *P*_bound_ ∝ *Z*_bound_. Because the data only provide information about the relative rate at which probes are selected, only the relative values of *α_a_* and *α_a:b_* are meaningful in this limit. Fourth, while PSAM models can be accurate for close-to-consensus sequences, they severely underestimate the affinity of far-from-consensus sequences, for which non-specific binding is dominant^56^. This can be addressed by including a non-specific binding term *α*_N.S._ in *Z*_bound_. Finally, it is sometimes important to include a factor *ω_a_*(*x*) that models biases in binding along the probe. Putting all of this together gives that the partition function in selection round *r* is given by:

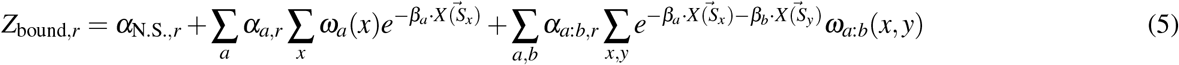

The binding probes typically feature a variable region flanked by constant sequences. The sliding window sum over subsequences *S_a_* can be configured to include *f_a_* letters from the flanking sequences. By default, the sum runs over both strands, but it can be restricted to only one strand (which is useful for modeling RNA and peptides).

#### Selection model

The selection model predicts the relative concentrations *f_i,r_* of each binding probe *i* each selection round *r*. By default, the concentrations in two subsequent rounds are related through an enrichment factor proportional to the binding. It is convenient to express this as

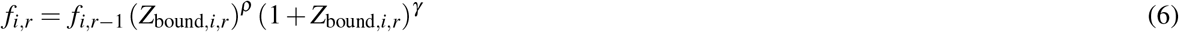

where *Z*_bound,*i,r*_ is the partition function evaluated for probe *i* in round *r*. Experiments conducted in the low–protein-concentration limit are modeled by setting (*ρ, γ*) = (1,0). Binding saturation can be accounted for by setting (*ρ, γ*) = (1, −1).

Some experiments (such as *K*_D_-seq, see below), do not use multiple rounds of binding enrichment and are better modeled using

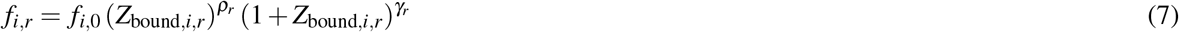

Finally, kinetic experiments that enrich and sequence modified or unmodified probes can be modeled using the constant-rate enrichment model:

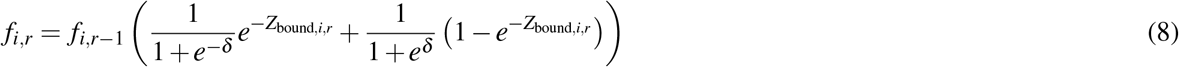

Here *δ* → ∞ and *δ* → −∞ correspond to the unmodified and modified fractions, respectively.

#### Sequencing model

The sequencing model computes the likelihood of the observed count tables *k_i,r_* given the relative concentrations *f_i,r_* predicted by the selection model. The counts are assumed to follow a Poisson distribution with expectation value

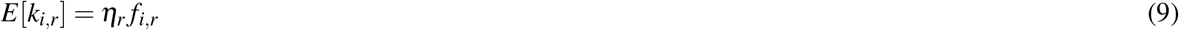

Here the parameter *η_r_* normalizes the relative probe concentration and adjusts to the correct sequencing depth. The (rescaled) likelihood is then

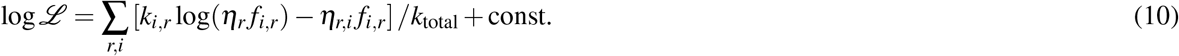

where *k*_total_ is the total number of reads and where the last term is independent of model parameters and can be ignored for the purpose of optimization. Because *f_i,r_* is proportional to *f*_*i*,0_, the latter parameter can be optimized analytically and substituted back into Eq. 10, giving

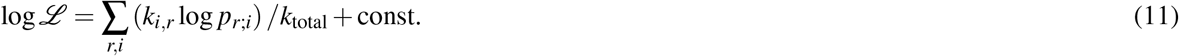

where 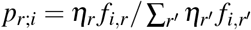. Note that Eq. 11 also can be derived by assuming the counts for each probe follow the multinomial distribution across columns with probability *p_r;i_*. Also note that because all unobserved probes have *k_i,r_* = 0 and do not contribute to the likelihood, the sum over *i* only runs over the the observed probes. This is a major advantage compared to NRLB^22^, where the sum is over all 4^*L*^ probes, with *L* is the number of variable positions. This sum can only be evaluated using dynamic programming and this restricts NRLB to data from only a single round of affinity-based enrichment in the absence of saturation. Finally, note that Eq. 11 is independent of the initial probe frequencies *f*_*i*,0_, meaning that initial library need not be random but can consist of genomic DNA fragment or custom-designed sequences.

#### Multi-experiment learning

ProBound simultaneously models multiple experiments by computing the likelihood 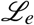 of each experiment *e* and then optimizing the combined likelihood

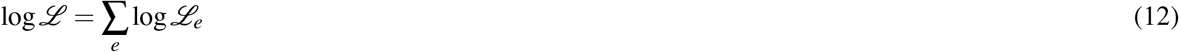

The precise way in which the likelihood 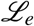 is evaluated can be tailored to the details of each experimental design:

1. A different configuration of binding modes and their interactions can be chosen for each experiment when computing *Z*_bound_ when desired.
2. The binding mode (and interaction) activities can either take independent values *α_a,e_* in each experiment or be constrained to *α_a,e_* = [P_*a*_]_*e*_*α_a_* where *α_a_* is the global activity of binding mode *a* and [P_*a*_] is a set parameter. The latter is useful when integrating experiments conducted at different protein concentrations, or in kinetic assays where [P_*a*_] is set to the treatment time.
3. Chemical modifications are encoded by expanding the alphabet and transliterating letters appropriate experiments. For example, meCpG modifications can be encoded using the alphabet `ACcGgT`, the complementarity rules `A ↔ T, C ↔ G, c ↔ g`, expanding the feature set Φ of the binding model to inlude the additional letters, and performing the transliteration `CG → cg` for methylated probes.

#### Regularization

Three regularization terms were included to avoid overfitting and to improve the stability of the numerical optimization: The first was a *L*_2_ regularization term for the parameter vector

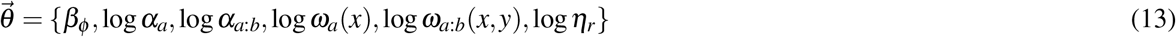

with weight *λ*. The second term was inspired by the Dirichlet distribution which commonly is used as a prior for probability parameters. For each feature *ϕ* thus we identified all features Φ^*c*^(*ϕ*) that are of the same class *c* (monomer, or dimer with the same spacing) and located at the same position within the binding site, and then define a feature probability

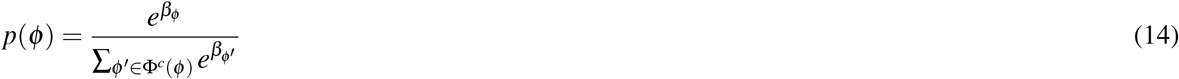

The regularization term is then computed as the rescaled log-PDF of *p*(*ϕ*) in the Dirichlet distribution

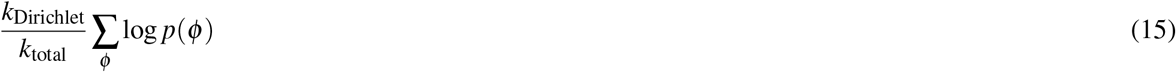

where *k*_Dirichlet_ is analogous to a pseudocount. The final regularization term in the likelihood is defined as

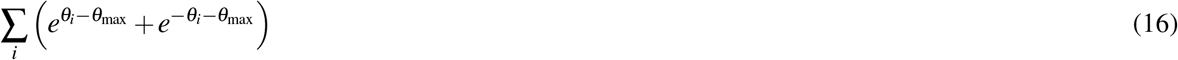

and introduces an exponential barrier (by default *θ*_max_ = 40) that prevents the optimizer from failing or getting trapped in regions with large numerical errors.

#### Procedure for setting k_Dirichlet_

The importance of the Dirichlet regularizer in Eq. 15 is set by *k*_Dirichlet_. For fits with all-by-all interactions, the inferred coefficients tended to be unstable for small values of *k*_Dirichlet_. While increasing *k*_Dirichlet_ stabilizes the coefficients, they shrink towards zero when *k*_Dirichlet_ is excessively large. We thus developed a procedure for setting *k*_Dirichlet_ and applied it uniformly in our analysis of Dll (Figure 4b), RBFOX2 (Figure S5j), and Src (Figure 6b). In this procedure, we ran ProBound using a wide range of Dirichlet weights (*k*_Dirichlet_ ∈ {0, 10, 20, 50, 100, 200, 500, 1000, 2000}), fixed the monomer coefficients 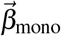 and dimer coefficients 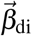 in each resulting model using the mismatch gauge (see below), and computed the pairwise Pearson correlation *r*^2^ between the inferred 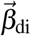 for different values of *k*_Dirichlet_. The resulting matrix *r*^2^(*k*_1_, *k*_2_), where *k*_1_ and *k*_2_ are values of *k*_Dirichlet_, had a block-like structure where 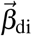 was highly correlated for large values of *k*_1_ and *k*_2_ but only weakly correlated when *k*_1_ or *k*_2_ was small. We considered the coefficients to have stabilized when *r*^2^ > 0.8 between a model and the model with the next-smaller value of *k*_Dirichlet_. Using this procedure, we fixed *k*_Dirichlet_ to be 200 for RBFOX2, 200 for the single-experiment Dll analyses, 1000 for the multi-experiment Dll analyses, and 50 for Src.

#### Model optimization scheme

ProBound optimizes the above model by first restricting it to only include the first binding mode (and non-specific binding) and optimizing this model, and then sequentially including and optimizing additional binding modes (and interactions as they become possible). As each new binding mode *a* (or interaction *a*: *b*) is included and optimized, the algorithm takes seven sub-steps: (i) heuristic adjustment of *α_a_* (or *α_a:b_*) so that it is expected to contribute to 5% to *Z*_bound_; (ii) freezing the values of all model parameters; (iii) unfreezing and optimizing ***η*** to avoid shocks from incorrectly predicted sequencing depth; (iv) unfreezing and optimizing the monomer features in 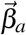 mode to give an initial binding model (***ω***_*a:b*_(*x,y* is unfrozen and optimized for interactions); (v) greedy exploration of alternative binding models with different frame shift (shifting the recognized sequence features to left or right), footprint (expanding the region of feature recognition to the left and/or right) or flank length (including subsequences located further into the fixed flanking regions when computing *Z*_bound_); (vi) sequential unfreezing and optimization of dimer features and ***ω***_*a*_(*x*) if applicable; (vii) unfreezing of all model parameters. At each substep, L-BFGS is used to optimize the unfrozen parameters. By default, the parameters are seeded with small random numbers, but the binding modes can also optionally be seeded using IUPAC codes. Additional constraints can be imposed on the parameters to implement reverse-complement symmetric binding modes or translationally symmetric interactions.

#### Gauge fixing

Models with pairwise letter interactions are over-parametrized, meaning that an infinite set of parameter values 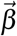 encode the same sequence specificity. Specifically, for any binding site sequence *S*, 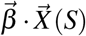 is invariant under transformations of the form

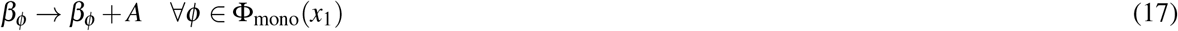

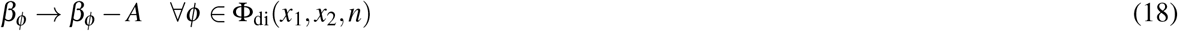

where Φ_mono_(*x*_1_) is the set of monomer features at position *x*_1_, Φ_di_(*x*_1_, *x*_2_, *n*) is the set of dimer features connecting positions *x*_1_ and *x*_2_ and with *n* at *x*_2_, and *A* is the transformation coordinate. For visualization and model comparison purposes, it is convenient to select one representative model for each sequence specificity (analogous to gauge fixing in physics). We here use a convention we call the ‘mismatch gauge’. In this convention, the coefficients are such that, first, only one monomer coefficient contributes for single-edit variations of reference sequence *S*_0_, and, second, at most one of the dimer coefficients contribute for each double-edit variations of *S*_0_. After imposing mutation gauge, the resulting PSAMs were visualized using standard energy logos^57^ and the interaction coefficients were displayed using heat maps.

### Benchmarking ProBound

#### How fits were trained, trimmed, and selected

To benchmark ProBound, we first curated a training database of published TF SELEX datasets^9,10,12,14,15,26–28^. Datasets with low sequencing depth or low enrichment were filtered out as described below. Each dataset was then analyzed by ProBound using three settings that differed in the number of binding modes and in how non-specific binding was modeled (see Supplemental Methods).

For each resulting fit, one binding mode typically captured the TF sequence specificity and the other typically absorbed platform-specific biases. To automatically identify the TF mode, we computed a heuristic quality score, which favors modes that both are important for the fit and have high specificity, and selected the mode with the top score. This score was 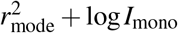, where 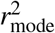 is the the Pearson correlation (across SELEX probes) of the log-transformed binding affinity predicted by the mode (plus an optimized non-specific term) and the log-transformed binding predicted by the full fit, and *I*_mono_ is the information content of the mononucleotide coefficients after imposing the mismatch gauge.

To automatically select which of the three settings produced the best fit in a way that does not give an unfair advantage when comparing to published models, we developed the quality score *S*_training_ which measures model performance in predicting the training data. *S*_training_ was defined to be the average of six sub-scores that quantify different aspects of model performance:

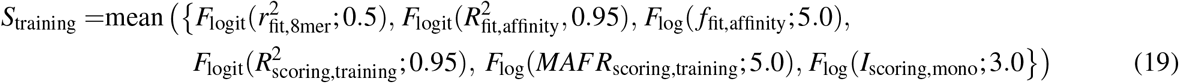

where the functions *F*_logit_(*x;x*_0_) = expit(logit(*x*) – logit(*x*_0_)) and *F*_log_(*x;x*_0_) = expit(log(*x*) – log(*x*_0_)) mapthemetric *x* to the unit interval such that the threshold *x*_0_ maps to 0.5. Here,

- 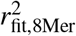 was computed by first using the full ProBound model to predict the training count table, then counting the number of occurrences 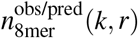 of each 8mer *k* in each round *r* of the of the observed and predicted count tables, then computing the observed and predicted 8mer enrichment between the first and last round using

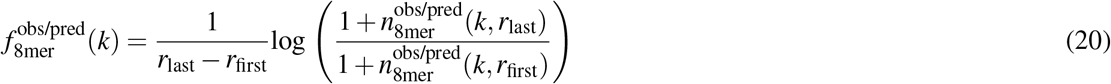

and finally computing the Pearson correlation between 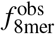 and 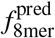.
- 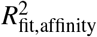 and *f*_fit,affinity_ were computed by first using the full ProBound model to predict the training count table. Then, for each pair of rounds subsequent rounds *r* and next(*r*) (ignoring rounds with less than 10,000 reads), the probes were sorted (conjointly in the observed and predicted tables) by the predicted enrichment between the rounds. The probes were then divided into bins *i* with associated the observed and predicted probe counts 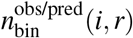 such that 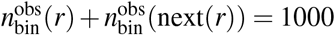 in each bin. After computing the observed and predicted enrichment using

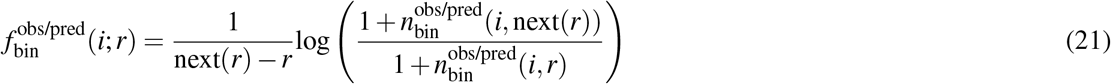

we finally computed the metrics

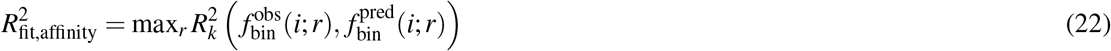

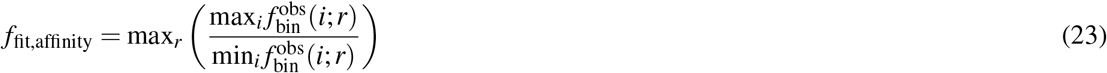

where 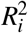 denotes the coefficient of variation evaluated across bins *i*.
- 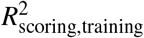 and *MAFR*_scoring,training_ were computed using the same method that was used to quantify generalization performance in predicting testing SELEX data (see below) but instead predicting the training data.
- *I*_scoring,mono_ is the information content of the scoring model, computed using the monomer coefficients after imposing the mismatch gauge.

#### Evaluation of Model Performance

To benchmark the resulting binding models, we curated a testing database of published SELEX (same as training database, but excluding the training dataset), PBM^58–60^ and ENCODE ChIP-seq^31^ datasets. We then quantified the ability of the above binding models to predict the testing data. Binding models and testing data were matched by TF and species; if no match was found, the matching criteria were expanded to consider orthologous human and mouse TFs. For comparison, we also downloaded binding models from the JASPAR, DeepBind, HOCOMOCO databases and the original HT-SELEX TF binding survey^26,30,32,33^ and repeated all analysis using these models.

For the SELEX and PBM experiments, we used the binding models to predict the total affinity (denoted *x_i_*) for each probe *i* and quantified how well these predictions agree with the measured binding *y_i_*. For the SELEX experiments, the signal consisted of the probe-count enrichment *k_i,r+1_/k_i,r_* between subsequent SELEX rounds (with maximum normalized to 1). For the PBM experiments, the background-subtracted and min-max normalized binding signal was used. For both platforms we encountered two challenges: First, the measurements for individual probes were too noisy to quantify model performance accuracy (for SELEX, typical sequences were observed just once; for PBM, the signal depends strongly on the position of the binding site in the probe, which varies). Inspired by earlier PBM analyses which removed position bias by considering the 8mer-binned median signal^29,61^, we sorted and binned the probes using *x_i_* (with bin size 500 for SELEX and 10 for PBM) and then computed the binned signal *y_i_* (using the bin-averaged enrichment, with pseudocount 1, for SELEX, and the median signal for PBM). Second, binding signals can be distorted by experimental artifacts such as binding saturation, background, and non-specific binding not modeled by the model. To correct for such distortions, *x_i_* was transformed using the binding saturation function:

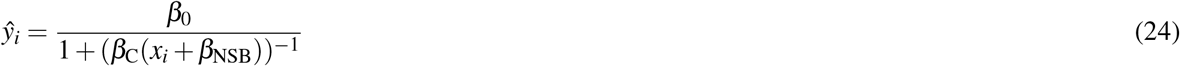

Here *β*_0_ sets the scale, *β*_C_ > 0 sets the concentration, and *β*_NSB_ sets the non-specific binding. These parameters were estimated by minimizing 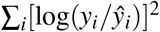 for SELEX (with *β*_0_ > 0 and *β*_NSB_ > 0) and 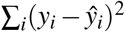 for PBM (for which *y_i_* can be negative). Model quality was then quantified using the coefficient of determination *R*^2^ of *y_i_* and *ŷ_i_* (on a logarithmic scale for SELEX) and the MAFR, which is defined as (max_*i*_*y_i_*)/*y*_bg_ where *y*_bg_ is the weakest signal detected by the model. To estimate *y*_bg_, we first defined a set of (binned) probes predicted to be bound as *ŷ_i_* > 1.25 *Q*_1_(*ŷ*) (where *Q*_1_ is the first quartile) and then defined *y*_bg_ to be the smallest value of *y_i_* identifying the bound set at 5% FDR. For multi-round SELEX experiments, *R*^2^ and the effective range were computed for all rounds and the largest values were recorded.

For the ChIP-seq experiments, we quantified model performance using the area under the precision-recall curve in classifying binding peak vs. background sequences. To get the peak sequences, we downloaded narrowPeak files from the ENCODE portal (see below) and extracted the genome sequence from the 500 peaks with the strongest enrichment. To generate the background set, we shifted the peak interval one peak length to the left and right and extracted the genome sequences.

#### Filtering of SELEX training datasets

We first curated a database of published SELEX experiments and downloaded the associated raw sequencing data^9,10,12,14,15,26–28^. Methylated SELEX experiments were not considered. For each experiment, we downsampled the sequencing libraries to contain at most 100,000 reads and tabulated the probe counts in each SELEX round. We then filtered out low-quality experiments using three criteria: First, low-coverage experiments were removed by requiring at least two rounds to have at least 10,000 reads. Second, experiments were discarded if no sequencing library before round three had 10,000 or more reads. Third, experiments with low-enrichment were discarded. The enrichment was quantified by first tabulating the frequencies *p*(*k, r*) (using pseudocount 5) of all 5mers *k* in each SELEX round *r*, and then, for each pair of rounds *r_i_* and *r_j_* with 10,000 or more reads, computing the rescaled KL divergence

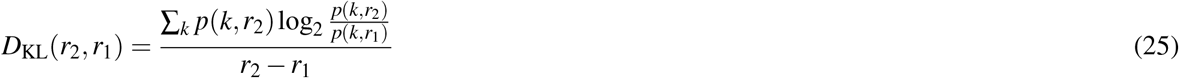

Only experiments with rescaled KL divergence exceeding 0.01 for at least one combination of rounds were retained.

#### Scoring of binding probes

In quantifying generalization performance, we predicted the occupancy of DNA sequences using both the ProBound binding models and previously published models. For DeepBind, we exponentiated the scores returned from the `deepbind` scoring tool, which is proportional to binding affinity. For JASPAR and original HT-SELEX TF survey, the binding models were position-frequency matrices (containing counts). These were first converted to position probability matrices (PPM, using a psuedocount of 1) which were then used to compute the binding probability at each offset in the sequence. The occupancy was then defined to be the sum of the binding probabilities. For HOCOMOCO, the binding models were PPMs and the occupancies were computed as described above.

#### ENCODE ChIP-seq datasets

ENCODE datasets were downloaded on December 2018 using the query string:

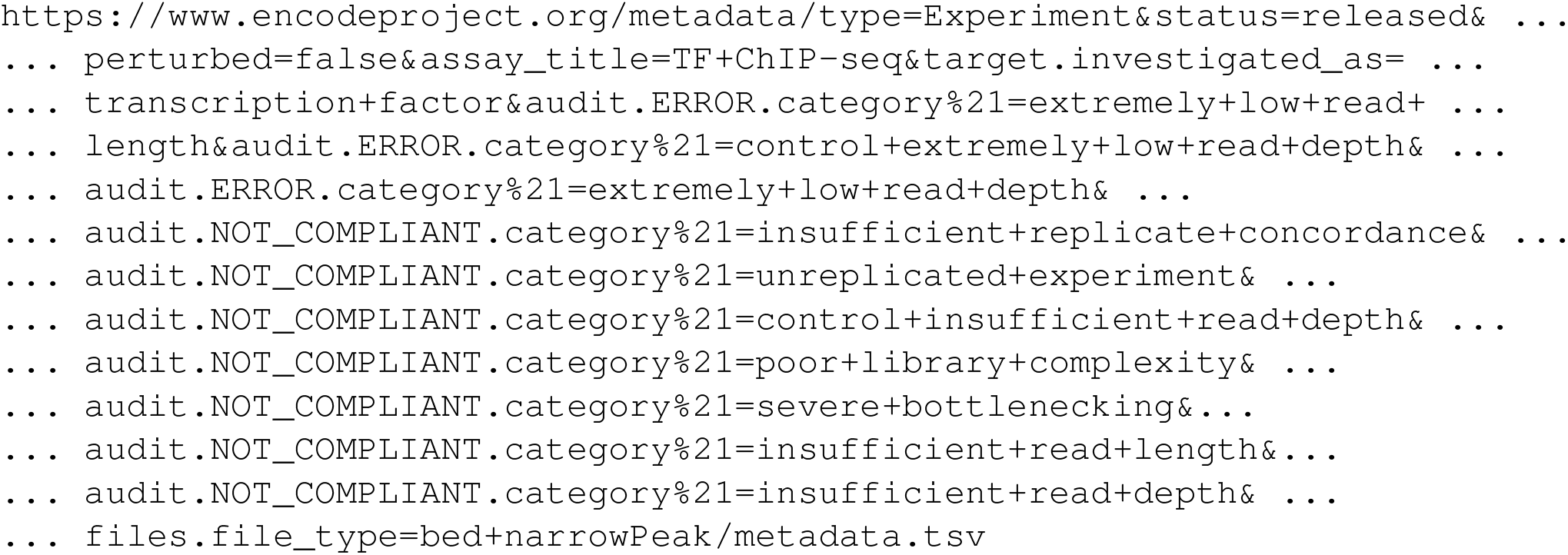

### Binding by multi-protein complexes

#### ProBound Analysis

ProBound was configured to jointly analyze SELEX experiments performed with different combinations of TFs, as described in the Supplemental Methods. In the case of Hth-Exd-Ubx, we analyzed published SELEX-seq experiments for Exd-Ubx, Hth, Exd, and Ubx. In addition, we preformed SELEX-seq for Hth-Exd-Ubx (see below). POU2F-GSC2 and ELK1-GCM1 were analyzed as described in the Supplemental Methods and Supplemental Table 4.

#### Experimental Protocol

The Hth-Exd-Ubx SELEX experiment was carried out following previously published methods^10,62^. Briefly, after expressing and purifying the wild-type homeodomain proteins, a final concentration of 50 nM was assembled, incubated with excess DNA (10-20 fold) for 30 minutes, and loaded onto an EMSA gel. A DNA library with 30 randomized bases was used. The TF-bound fraction was isolated from the gel, amplified, and either subjected to another round of enrichment or prepared for sequencing. Three rounds of enrichment were performed. After each selection round, the DNA was extracted from the gel and amplified by using Ilumina’s small RNA primer sets. Sequencing barcodes were added in a five cycle PCR step and the final library was gel-purified using a native TBE-gel before sequencing. Libraries were sequenced at the New York Genome Center using separate lanes on an Illumina HiSeq 2000 sequencing machine.

### Effect of DNA Methylation

#### ProBound Analysis

ProBound learns methylation-aware binding models by jointly analyzing normal and methylated SELEX libraries after encoding the methylation state of each basepair using an extended alphabet (see Figure S3a and configuration in Supplemental Methods). Encoding methylation status in this manner allows us to infer the position-specific free energy impact of such chemical modifications. For the ATF4/CEBP*γ* homo- and hetero-dimers, we jointly analyzed two published EpiSELEX-seq experiments for ATF4 and CEBP*γ*, and a new EpiSELEX-seq experiment that included both ATF4 and CEBP*γ*. We also generated EpiSELEX-seq data for CEBP*γ* in combination with the chemical modifications meCpG, 5hmC, and 6mA.

#### Experimental Protocol

ATF4 protein purification and EpiSELEX-seq experiments were performed as described previously^15^. Purified CEBP*γ* protein was kindly donated by the Lomvardas lab at the Zuckerman Institute at Columbia University. To generate randomized 5hmC or 6mA libraries, single-stranded oligos with a 16-bp randomized region were ordered from TriLink Biotechnologies, substituting i) deoxycytidine triphosphate (dCTP) with deoxy-(5hm)-cytidine triphosphate (d5hmCTP), or ii) deoxyadenosine triphosphate (dATP) with deoxy-(6m)-adenosine triphosphate (d6ATP) during the synthesis step. For double-stranding, a standard mix of deoxy-nucleotides was used, resulting in hemi-modified libraries. meCpG libraries were generated by enzymatic treatment with M.SssI (NEB) as described previously^15^. The library sequences consisted of left and right constant adapters (GGTAGTGGAGG- and -CCAGGGAGGTGGAGTAGG respectively) flanking a library specific barcode and a 16bp randomized sequence:

- no modification: -TGGG-CCTGG-N16-
- meCpG: -GCAC-CCTGG-N16-
- 5hmC-Library: -CAGT-CCTGG-N16- (5hmC instead of C in `16N`)
- 6mA-Library: -AGTG-CCTGG-N16- (6mA instead of A in `16N`)

#### GLM analysis of ATF4 and CEBPγ ChIP data

To estimate the effect of DNA methylation on *in vivo* AFT4 and CEBP*γ* binding, we first scanned the genome for close-to-consensus motif matches *i* with `CG` at positions predicted by the model to have strong methylation readout: TGA<u>CG</u>TCA and TGA<u>CG</u>T<u>CG</u> for ATF4:AFT4; TTG<u>CG</u>CAA for CEBP*γ*:CEBP*γ*; and TTG<u>CG</u>TCA and TTGCAT<u>CG</u> for CEBP*γ*:ATF4. We next downloaded aligned ATF4 and CEBP*γ* ChIP-seq reads and matched input from ENCODE (ENCFF872NFM, ENCFF801LQC, ENCFF713PVH), extended the alignments to 125bps, and computed the genome coverages (*k*_ATF4,*i*_, *k*_CEBp*γ,i*_, *k*_input,*i*_) at each motif match. The DNase-seq coverage (*k*_DNase_,_i_, ENCFF971AHO) and bisulfite sequencing methylation status (*f*_meCpG,*i*_, ENCSR765JPC, binarized using 20% and 80% thresholds, and keeping matches with at least 10 reads) were also recorded. We finally modeled the ATF4 and CEBP*γ* ChIP-seq coverage at the relevant motif matches (excluding CEBP*γ*:CEBP*γ* matches for ATF4 and ATF4:ATF4 matches for CEBP*γ*) using two separate binomial generalized linear models:

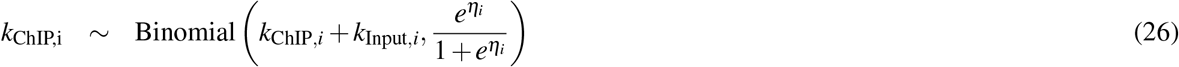

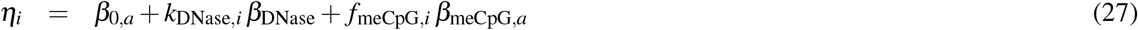

In this model, *β*_0,*a*_ encodes the relative affinity of motif *a*, *β*_DNase_ encodes the impact of DNA accessibility, and *β*_meCpG_ encodes the impact of DNA methylation for motif *a* and is the sought-after variable. The significance of the methylation readout was assessed using a F-test (see Supplemental Table 2). For TGA<u>CG</u>T<u>CG</u>, we assumed that the methylation readout of the two CGs contribute independently and that the readout of the central CG can be estimated using the sequence TGA<u>CG</u>TCA.

### Inferring Absolute K_D_’s

The *K*_D_-seq assay incubates a protein TF (or other protein) with a library of DNA probes (or RNA or peptide probes), separates the bound and free probes, and sequences the input (I), bound (B) and free (F) fractions. In equilibrium, the probability that probe *i* is bound or free is given by

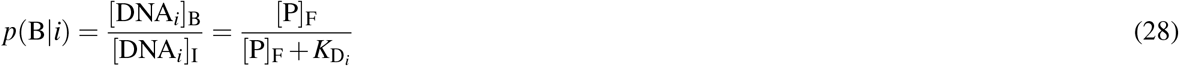

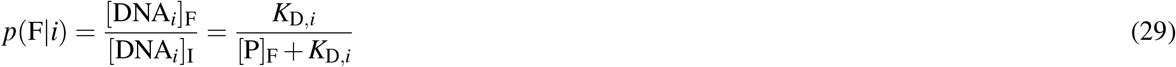

where [DNA_i_]_I_, [DNA_i_]_B_, and [DNA_i_]_F_ are the probe concentrations in the input, free and bound libraries, [P]_F_ is the free protein concentration, and *K*_D,*i*_ is the dissociation constant that we wish to measure. The sequencer does not measure *p*(B|*i*) or *p*(F|*i*) directly but rather gives the probe counts *k*_*i*,I_, *k*_*i*,B_, and *k*_*i*,F_. The expectation values of these counts are given by

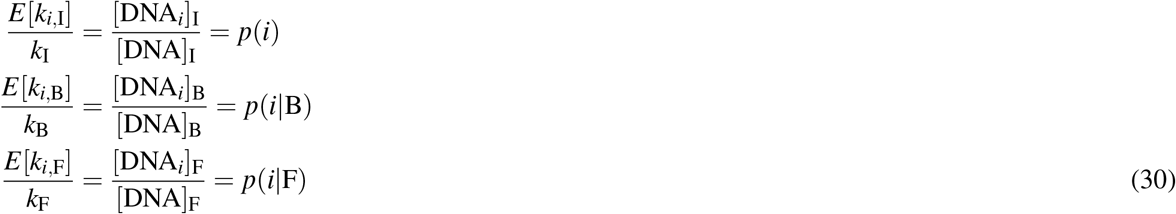

where [DNA]_I_, [DNA]_B_, [DNA]_F_ are the DNA concentrations in the in the respective fractions, *k*_I_, *k*_B_ and *k*_F_ are the sequencing depths of the libraries which are treated as fixed experimental setting. To estimate the dissociation constants, note that

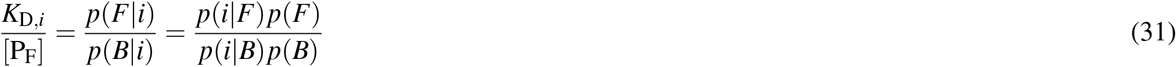

where *p*(*B*) and *p*(*F*) are the net fractions of DNA that is bound and free. Intuitively, these can fractions can be estimated from the data by finding the values that make the observed probabilities in Eq. 30 satisfy the sum rule:

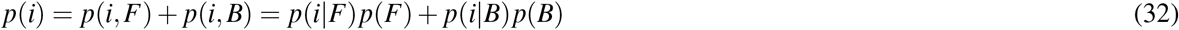

ProBound can be configured to learn a *K_D_* model by analyzing the probe frequencies in the input, bound and free libraries (*r* = {I, B, F}). Specifically, configuring ProBound to use the non-cumulative enrichment model (Eq. 7) with *ρ_r_* = {0,1,0} and *γ_r_* = {0, −1, −1} and restricting the activities to be constant across columns implements the binding probabilities in Eq. 29. With these settings,

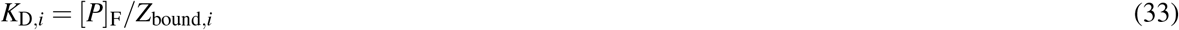

The ProBound model implicitly encodes p(B); this value can be found by equating the expected counts in ProBound

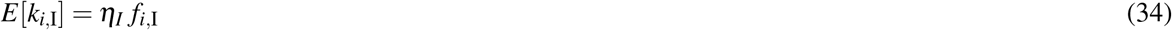

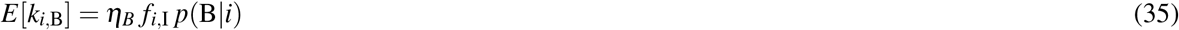

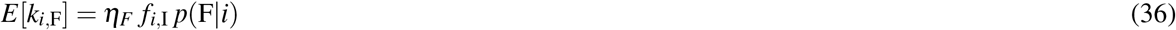

with the corresponding expectation values in Eq. 30, computing the bound-to-input ratio, and using Bayes’ theorem to simplify, giving

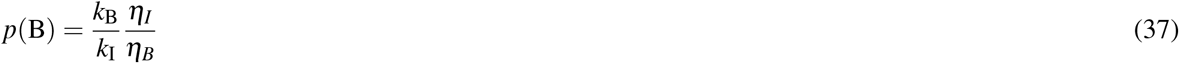

Test the modeling assumptions (c.f. Figure 4c), the probes were binned by the predicted *K*_D,*i*_, and, for each bin, the observed and predicted binding probabilities were computed using

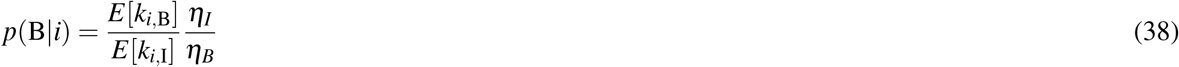

Here *E*[*k*_*i*,B_] and *E*[*k*_*i*,I_] were evaluated using the observed and predicted read counts in each bin.

#### Experimental Protocol

6xHis tagged Drosophila Distalless (Dll) protein lacking amino acids N terminal to its homeodomain (DllΔN) was purified by standard procedures. 0.05% Tween-20 was included in the lysis buffer and in the elution buffer to prevent the target protein from sticking to plasticware. The purified protein was quantified by Bradford assay, using BSA as the standard. The 10mer R0 library was generated by annealing the library oligo (GTTCAGAGTTCTACAGTCCGACCTGG −10N -CCAGGACTCGGACCTGGACTAGG) and the SELEX-R primer (CCTAGTCCAGGTCCGAGT), followed by a Klenow mediated primer extension reaction. The library DNA was purified using Qiagen minElute columns, and were quantified using nanodrop. The SELEX procedure was largely the same as previously described^10^, except that a Cy5 labeled DNA probe, instead of a P32 labeled probe, was used as the marker to indicate where the bound and unbound fractions were. The Cy5 labeled DNA probe was generated by annealing a Cy5 labeled primer to a DNA probe with the desired DNA sequence, followed by Klenow reaction. EDTA was used to stop the reaction. The probe was directly used in the binding reaction, without further purification.

For each SELEX condition, 15*μ*l of protein solution (at 2x final concentration) in dialysis buffer (20mM HEPES pH8.0, 200mM NaCl, 10% glycerol, 2mM MgCl_2_, 0.05% Tween-20) was made. The library mixture was made by adding desired amount of the R0 library to 6*μ*l of 5x binding buffer (50mM Tris-HCl pH7.5, 250mM NaCl, 5mM MgCl_2_, 20% glycerol, 2.5mM DTT, 2.5mM EDTA, 125ng/*μ*l polydIdC, 100ng/*μ*l BSA, 0.125% Tween-20), and filling to 15*μ*l with H_2_O. The protein and DNA parts were mixed and incubated at room temperature for 30 to 40 minutes before loading the gel. For Cy5 labeled markers, 15*μ*l of 200nM DllΔN in dialysis buffer was mixed to 15*μ*l of DNA mixture (6*μ*l 5x binding buffer, 8*μ*l H_2_O and 1*μ*l 200nM probe), and was incubated at room temperature for 30 to 40 minutes.

After running the gel, gel slices corresponding to the bound and unbound fractions were cut from the gel, and were each place in a 500*μ*l tube with several needle poked holes at the bottom. The 500*μ*l tubes were each placed within a 2ml tube, and was spun at max speed at room temperature to smash the gel. 650*μ*l of DNA extraction buffer (10mM Tris-HCl, pH7.5, 150mM NaCl, 1mM MgCl_2_, 0.5mM EDTA, pH 8.0), and 50*μ*l of 20% SDS were added to each smashed gel sample, and the tubes were rotated at room temperature for 2 to 4 hours. The tubes were then spun at max speed at room temperature for 2 minutes. 650*μ*l of sample was transferred to a Spin-X filter column, and was spun at room temperature at the max speed for 2 minutes. The DNA in flow through was purified by phenol chloroform extraction followed by isopropanol precipitation. 20*μ*g of glycogen was used to facilitate precipitation, and the DNA pellet was dissolved in 20*μ*l of Qiagen EB buffer.

Each purified SELEX DNA was properly diluted such that the following PCR program gave good library yield for all samples. The 1-step library preparation was done in a 50*μ*l reaction, which contains 5*μ*l of properly diluted SELEX DNA, 10nM of one of the 8 SELEX-for primers, 10nM of the common SELEX-rev primer, 1 *μ*M of NEB universal primer for Illumina, and 1 *μ*M of selected NEB index primer for Illumina. PCR was done with the Phusion DNA polymerase (NEB), using the following program: 1 cycle of 98°C for 30 seconds; 5 cycles of 98°C for 10 seconds, 60°C for 30 seconds, and 72°C for 15 seconds; 10 cycles of 98°C for 10 seconds, and 65°C for 75 seconds; 1 cycle of 65°C for 5 minutes; and hold at 4°C. Amplified libraries were purified using 1.5 volume (75*μ*l) of Ampure beads, and eluted with 15*μ*l of Qiagen EB buffer. The libraries were pooled and sequenced using Illumina Nextseq 550, following standard procedures. The forward primers consisted of consisted of left and right constant sequences (ACACTCTTTCCCTACACGACGCTCTTCCGATCT- and -GTTCAGAGTTCTACAGTCCGA repectively), flanking a library specific barcode: 1) `--`, 2) `-AGAC-`, 3) `-TCAGAC-`, 4) `-CAGAC-`, 5) `-C-`, 6) `-GAC-`, 7) `-AC-`, and 8) `-TTCAGAC-`. In addition we used the reverse primer GACTGGAGTTCAGACGTGTGCTCTTCCGATCT-CCTAGTCCAGGTCCGAGT, the NEB universal primer AATGATACGGCGACCACCGAGATCTACACTCTTTCCCTA-CACGACGCTCTTCCGATCT, the NEB index primer `CAAGCAGAAGACGGCATACGAGAT-[6bp index]-GTGACTGGAGTTCAGACGTGTGCTCTTCCGATCT`.

#### EMSA validation

The same batch of the DllΔN protein that was used in the SELEX experiments was also used in the measurement of the absolute *K_D_* values of DllΔN to selected DNA sequences. The EMSA experiments were performed following regular protocol. Briefly, the protein was diluted with dialysis buffer to 2x of the desired final concentration in a total volume of 15*μ*l. The DNA mixture was made by mixing 6*μ*l of 5x binding buffer, 8*μ*l of H_2_O, and 1*μ*l of 200nM Cy5-labeled DNA probe. The DNA probes had the same flanks as the 10mer SELEX library, and the indicated middle 10bp. The protein part and the DNA part were mixed well, and incubated at room temperature for 30 to 40 minutes before loading the 0.5X native TBE gel.

After running the gel, an image was taken using the Typhoon imager, and the band intensity was quantified using FIJI v1.52n (see Supplemental Table 5). Briefly, each band was selected using the rectangle selection tool, and the selected regions were converted to histograms. A straight line was drawn at the bottom of each histogram, and the areas of the enclosed peak regions were quantified and used as band intensity.

*K*_D_ was finally estimated used non-linear binding curve fitting. The intensity of the bound band decreased with migration distance (data not shown). We therefore estimated the fraction of bound probes as *y*_Bound_/(*y*_Bound_ + *α y*_Free_), where *y*_Bound_ and *y*_Free_, respectively, are the intensities of the bound and free band, and *α* corrects for the migration-induced signal loss. In equilibrium, the predicted bound fraction equals

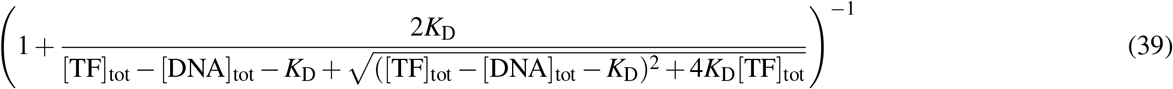

where [TF]_tot_ and [DNA]_tot_ are the total TF and DNA concentrations, respectively. For each probe, *K*_D_ and *α* were estimated by minimizing the squared difference between the estimated and predicted bound fractions across all DllΔN concentrations.

### Peak-free motif discovery from ChIP-seq data

To analyze the GR ChIP-seq data from the IMR90 cell line^47^, we first aligned the (single-end) Input and ChIP reads to the genome and extracted a sufficiently long (200bp) sequence downstream of the 5’-end genomic position of the mapped read. Next, we randomly sampled 10^6^ reads from each library and constructed a count table containing the Input and ChIP read counts in the first and second columns, respectively. ProBound was then configured to model this table as a single-round SELEX experiment. Because GR binds DNA as a homodimer, we configured ProBound to impose reverse-complement symmetry while fitting free-energy parameters the primary motif. We then iteratively added three additional binding modes to the model to capture the influence of potential co-factors. To analyze the GR ChIP-seq data from the murine hippocampus^48^, we followed a similar procedure and constructed one count table for each of the three CORT concentrations (sampling 10^5^ sequences per library) and then configured ProBound to jointly model all count tables using a single reverse-complement-symmetric binding mode.

### Tyrosine kinase sequence recognition

#### ProBound Analysis

In this assay, a library of peptide substrates *S_i_* is treated with a enzyme *E* and the concentrations of the products *P_i_* is quantified using high-throughput sequencing (see below). This reaction can be modeled using Michaelis-Menten kinetics generalized to multiple substrates:

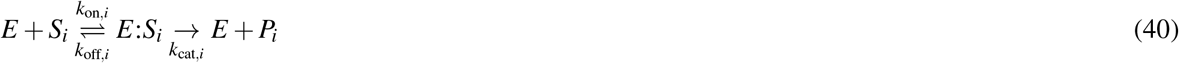

In the limit of low enzyme concentration, the reaction quickly reaches a quasi-steady state with

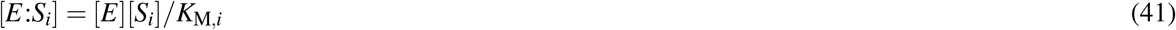

where *K*_M,*i*_ = (*k*_off_ + *k*_cat,*i*_)/*k*_on,*i*_ is the Michaelis constant for substrate *i*. In this limit, the change in substrate concentration is given by

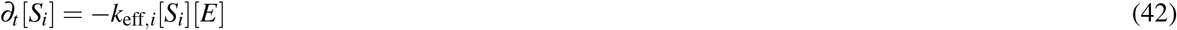

where *k*_eff,*i*_ = *k*_cat,*i*_/*K*_M,*i*_ is the catalytic efficiency. Integrating this equation yields

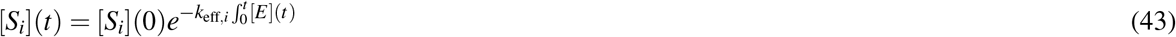

where [*S_i_*](0) is the substrate concentration right after the quasi-equilibrium was reached. The concentrations in the product library can then be expressed as

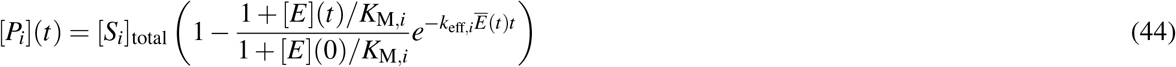

where [*S_i_*]_total_ = [*S_i_*] + [*E*:*S_i_*] + [*P_i_*] is concentration in the initial library and 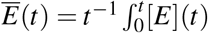 is the time-averaged enzyme concentration. This can be simplified further by noting that only a small fraction of substrates are bound in the limit of low enzyme concentration

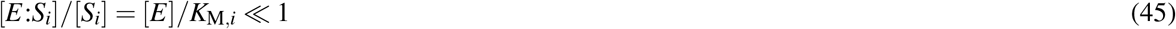

and thus

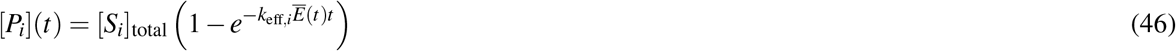

Note that the selection only differs between probes through *k*_eff,*i*_. ProBound can thus model the assay using Eq. 8 with *δ* → −∞ and

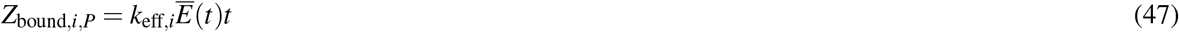

Here 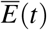 depends on both *K*_D,*i*_ and [*S_i_*] throughout the reaction and is generally unknown. We here assume that most enzyme is free so that 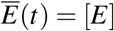; a lower (free) enzyme concentration would lead to a global rescaling of *k*_eff,*i*_ but not affect the relative efficiency or its sequence dependence.

#### Preparation of degenerate peptide library to profile tyrosine kinase specificity

The degenerate peptide library contained 11-residue sequences with five randomized amino acids flanking either side of a fixed central tyrosine residue. These sequences were fused to the eCPX bacterial surface display scaffold^63^. To clone this library, we first amplified the eCPX-coding sequence with a 3/ Sfil restriction site. This was fused to the random library in another PCR step using the following degenerate oligonucleotide: `GCTGGCCAGTCTGGCCAG-NNSNNSNNSNNSNNStatNNSNNSNNSNNSNNS-GGAGGGCAGTCTGGGCAGTCTG`, which contains a 5’ Sfil site. The resulting amplified product was digested with SfiI restriction endonuclease, purified, and ligated into the SfiI-digested pBAD33-eCPX plasmid, as described previously^53^. The ligation reaction was concentrated and desalted, then used to transform DH5*α* cells by electroporation. Transformed cells were grown overnight in liquid culture, then the plasmid DNA library was extracted and purified using a commercial midiprep kit.

#### Preparation of biotinylated antibody

The phosphotyrosine monoclonal antibody, pY20, conjugated to the fluorophore, perCP-eFluor 710 (Invitrogen, catalog 46-5001-42), was desthiobiotinylated before use in the specificity screen. The antibody was first purified away from bovine serum albumin (BSA) and gelatin by anion exchange using a salt gradient of 0 to 1 M NaCl in 0.1 M potassium phosphate buffer. The fractions that eluted after 0.2 M NaCl were pooled and then buffer-exchanged into 0.1 M potassium phosphate by dilution and centrifugal filtration. The antibody was then labeled in a 200 *μ*L small-scale reaction using the DSB-X labeling kit (Molecular Probes) according to the manufacturer’s instructions. Concentration of the antibody was monitored by its absorbance at 490 nm to determine percentage yield. The average final concentration of the antibody was around 0.2 mg/mL. The specificity of the antibody was validated using cells expressing displayed peptides. Cells treated with a tyrosine kinase without ATP show no background antibody staining. By contrast, cells expressing displayed peptides, treated with tyrosine kinase and 1mM ATP show increasing antibody staining as a function of phosphorylation time.

#### High-throughput specificity screen

The catalytic domain of the human tyrosine kinase c-Src was screened against the degenerate peptide library as described previously^53^, one main difference being the use of magnetic beads to isolate phosphorylated cells rather than fluorescence-activated cell sorting. In short, *E. coli* MC1061 cells transformed with the library were grown to an optical density of 0.5 at 600 nm. Expression of the surface-displayed peptides was induced with 0.4% arabinose for 4 hours at 25 °C. After expression, the cell pellets were collected and subject to a wash in phosphate buffered saline (PBS). Phosphorylation reactions of the library were conducted with 500 nM of purified c-Src and 1 mM ATP in a buffer containing 50 mM Tris, pH 7.5, 150 mM NaCl, 5 mM MgCl_2_, 1 mM TCEP, and 2 mM sodium orthovanadate. Time points were taken at 5, 20, and 60 minutes. Kinase activity was quenched with 25 mM EDTA and the cells were washed with PBS. Kinase-treated cells were labeled with roughly 0.05 mg/mL of the biotinylated pY20 antibody for an hour and then washed again with PBS containing 0.2% BSA.

The phosphorylated cells were isolated with Dynabeads^®^ FlowComp Flexi (Invitrogen) following the man-ufacturer’s protocol. In total, two populations were collected for each time point: cells that did not bind to the magnetic beads and eluted after each wash (unbound) and cells that bound to the magnetic beads and eluted after the addition of the release buffer (bound). After isolation of these two populations, the cell pellet was collected, resuspended in water, and then lysed by boiling at 100 °C for 10 minutes. The supernatant from this lysate was then used as a template in a 50 *μ*L PCR reaction to amplify the peptide-codon DNA sequence using the same forward and reverse TruSeq-eCPX primers as described previously^53^. The product of this PCR reaction was then used as a template for a second PCR reaction to append a unique 5’ and 3’ indices. The resulting PCR products were purified by gel extraction, and the concentration of each sample was determined using QuantiFluor^®^ dsDNA System (Promega). Each sample was pooled to equal molarity and sequenced by paired-end Illumina sequencing on a MiSeq instrument. The deep sequencing data were processed as described previously^53,64^. The paired-end reads were merged using FLASH^65^ and the adapter sequences were trimmed using the software Cutadapt^66^. The remaining sequences were translated into amino acid codes, and sequences containing stop codons were removed.

#### Validation measurement of phosphorylation rates

To validate predictions made by Probound, phosphorylation rates were determined *in vitro* using purified c-Src and 11 synthetic peptides (purchased from Synpeptide). The phosphorylation reactions were carried out at 37°C using 500 nM purified c-Src and 100 *μ*M peptide in a buffer containing 50 mM Tris, pH 7.5, 150 mM NaCl, 5 mM MgCl_2_, 1 mM TCEP, and 2 mM sodium orthovanadate. Reactions were initiated by the addition of 1 mM ATP, and at various time points, 100 *μ*L of the solution was quenched with 25 mM EDTA (every 10s for the faster reactions, every 2-10m for the slower reactions). Each reaction was carried out in triplicate.

The concentration of the substrate and the phosphorylated product at each time point was determined by reversed-phase HPLC with UV detection at 214 nm (Agilent 1260 Infinity II). A 40 *μ*L volume of the quenched reaction was injected onto a C18 column (ZORBAX 300SB-C18, 5*μ*m, 4.6 x 150 mm). A gradient system was used with solvent A (water and 0.1% TFA) and solvent B (acetonitrile and 0.1% TFA). Elution of the peptides was performed at flow rate of 1 mL/min using the following gradient: 0-2 min: 5% B, 2-12 min: 5-95% B, 12-13 min: 95% B, 13-14 min: 95-5% B, and 14-17 min: 5% B. The peak areas of the substrate and product were calculated using the Agilent OpenLAB software. The initial rate for each peptide was obtained by fitting a straight line to a graph of peak area as a function of time in the linear regime of the reaction progress curve and calculating the slope of the line.

## Acknowledgements

Research reported in this publication was supported by NIMH award R01MH106842 and NHGRI award R01HG003008 to H.J.B. and NIGMS award R35GM118336 to R.S.M. The content is solely the responsibility of the authors and does not necessarily represent the official views of the National Institutes of Health. We are grateful to John Hunt for valuable discussions about experimental methods for measuring dissociation constants.

## Author contributions statement

H.T.R. and H.J.B. developed the methodology with significant contributions from C.R. H.T.R. implemented ProBound with contributions from C.R., B.V.D. and H.H.A. S.F. performed the KD-seq experiments and validation measurements under supervision of R.S.M. J.F.K. performed the SELEX-seq and EpiSELEX-seq experiments and developed the GLM analysis under supervision of R.S.M. and H.J.B. A.L. performed the Src sequencing and validation experiments under supervision of N.H.S. B.B. developed the web portal under supervision of H.J.S., H.T.R. and C.R. X.L. performed the ASB ChIP-seq analysis. L.A.N.M and H.T.R. performed GR ChIP-seq ProBound analysis. H.T.R., C.R. and H.J.B. wrote the manuscript with input from all authors.

## Code availability

TF binding models and software for utilizing them can be accessed at motifcentral.org. The ProBound software can be run on a dedicated compute server located at probound.bussemakerlab.org.

## Data availability

The sequencing data generated during the current study have been deposited in the Gene Expression Omnibus (GEO, accession number GSE175942). Source data for Figs 4d and 6d have been provided in Supplemental Table 3 and 5.

---

**Figure S1:**
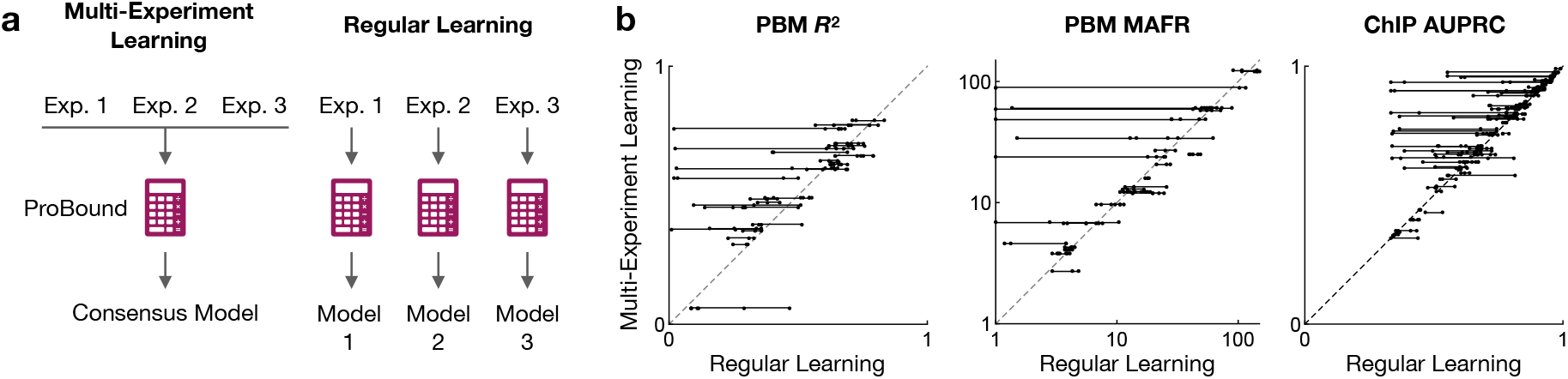
Integrative analysis of multiple TF SELEX datasets produces consensus binding models. **(a)** Schematic contrasting ProBound’s multi-experiment learning strategy that builds a consensus model for a TF by simultaneously training on all relevant SELEX data for the TF with the traditional approach that builds independent models for every individual dataset. **(b)** Generalization performance of consensus binding models (y-axis) and single-experiment models (x-axis) on three different metrics (scatterplots). Points correspond to models trained on individual experiments and lines connect experiments used to build the corresponding consensus model. Points above the diagonal correspond to instances where the consensus model outperforms single-experiment models.

**Figure S2:**
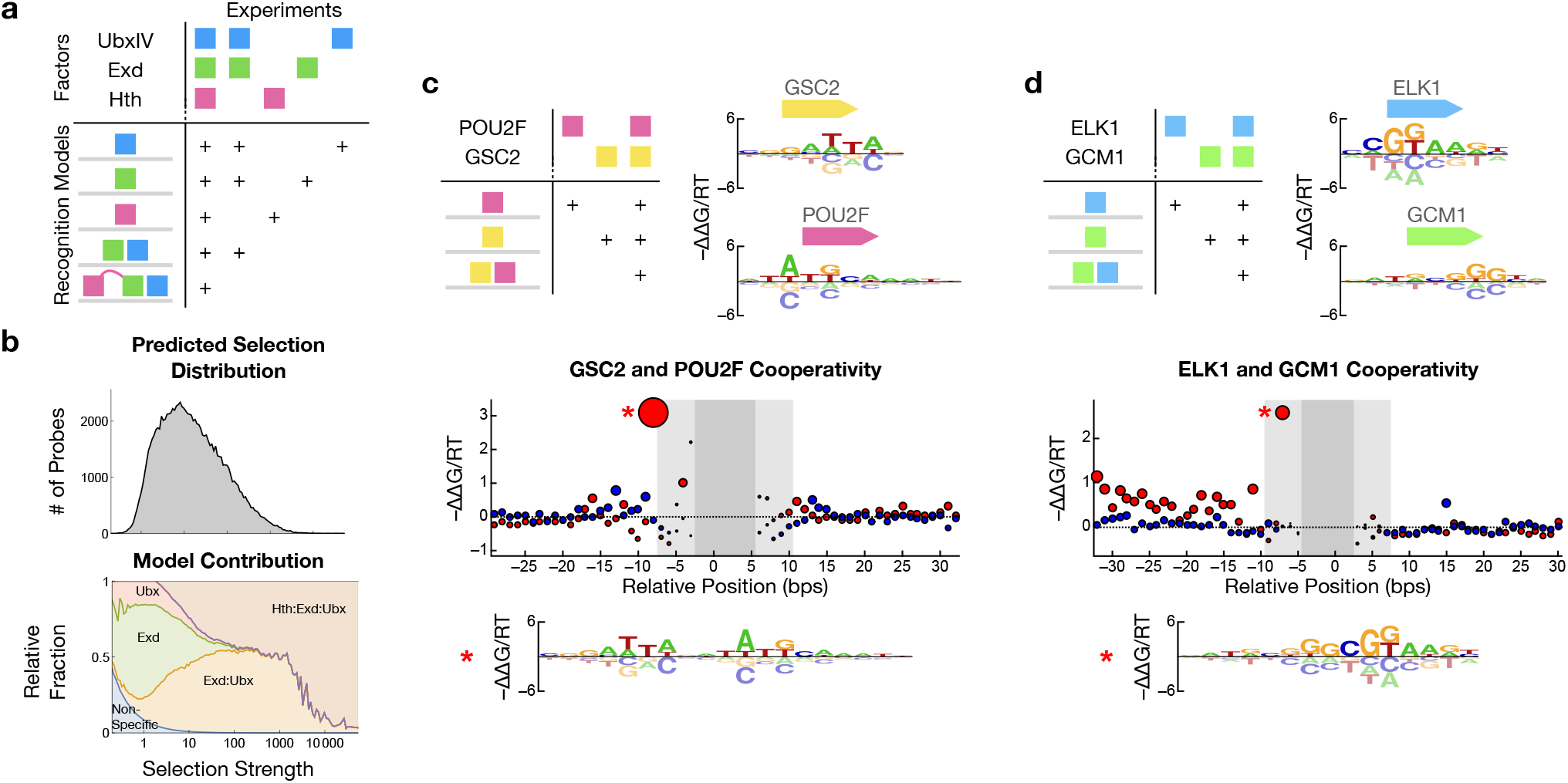
Integrative modeling to quantify TF binding cooperativity. **(a)** Schematic table describing the combinations of TFs assayed in five experiments (top) that were jointly analyzed to produce recognition models of the different monomers and their complexes (bottom) by explicitly defining which models can form in each experiment (+ sign). **(b)** Distribution of probes (top) and the predicted relative contribution of**l**every recognition mode (bottom) as a function of predicted binding selection strength (x-axis) in the first round of selection from SELEX-seq data assaying Hth, Exd, and UbxIV. (**c**) Integrative modeling of HT-SELEX and CAP-SELEX data for POU2F and GSC2 (schematic table) yields recognition models for the monomers (motifs) and binding cooperativity for GSC2:POU2F (scatterplot) as a function of relative position (x-axis) and orientation (red: parallel; blue: antiparallel). Motif (below) shows the configuration indicated on the plot. (**d**) Same as (c), except for the factors ELK1 and GCM1.

**Figure S3:**
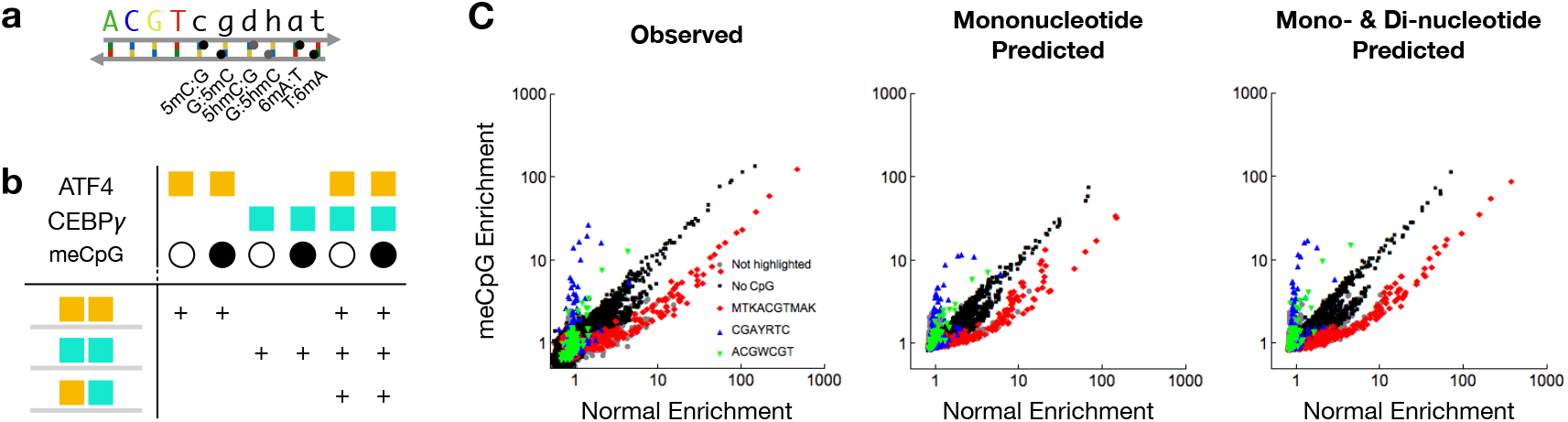
Learning methylation-aware binding models from EpiSELEX-seq data. (**a**) Alphabet used to represent normal and methylated base pairs. (**b**) Same as Figure S2a, but showing the combinations of ATF4, CEBP*γ*, and normal and methylated DNA that were included in each experiment and the resulting complexes that were modeled. (**c**) K-mer enrichment analysis for the observed ATF4 EpiSELEX-seq read counts (left), the counts predicted by a mononucleotide-only model (middle), and the counts predicted by a mono-and di-nucleotide model (bottom). Each scatterplot compares the 8mer enrichment observed in the normal (x-axis) and methylated (y-axis) libraries. Every point represents an 8mer and is colored according to the legend; color is assigned based on a 6bp matching substring between the 8mer and the IUPAC code.

**Figure S4:**
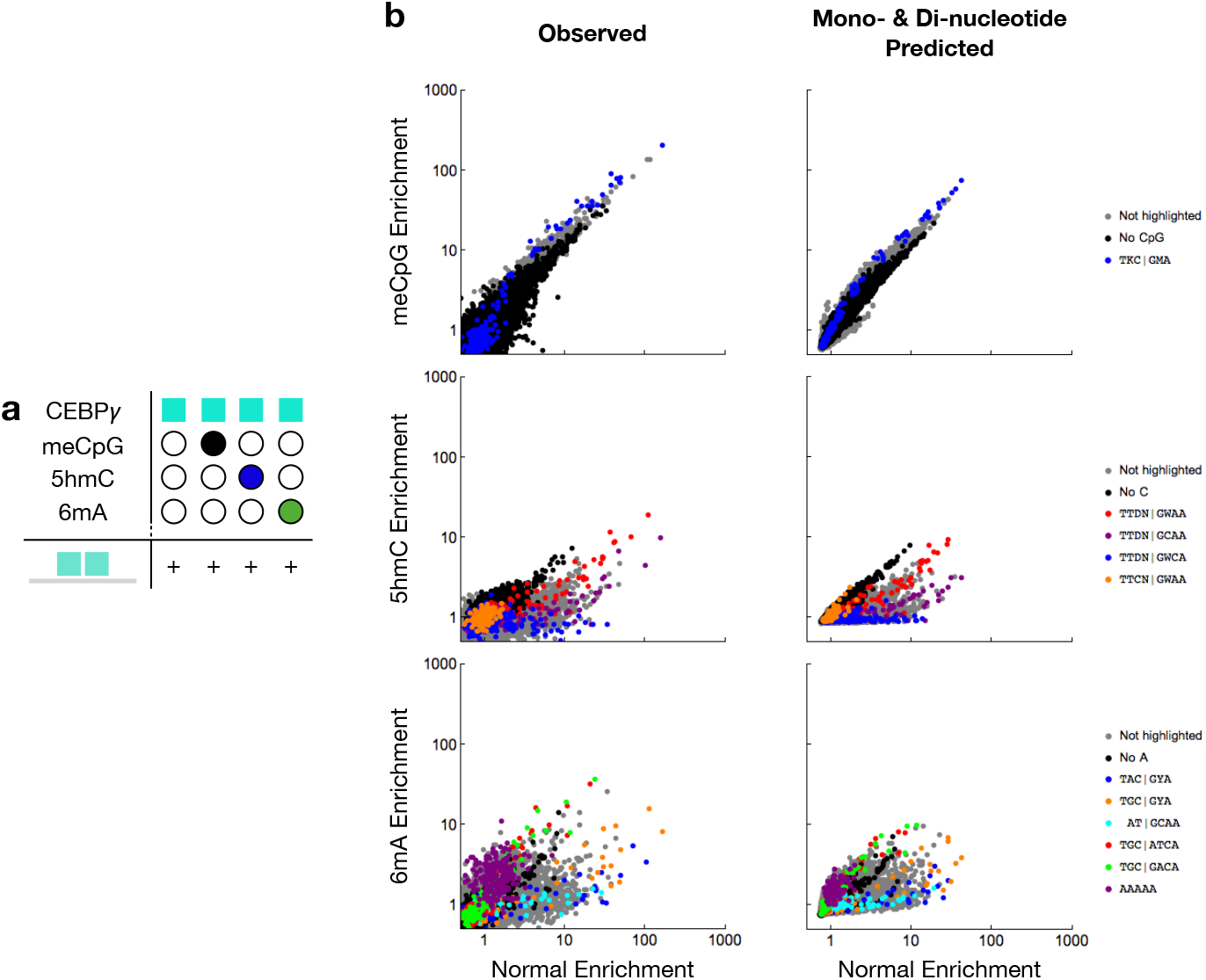
Extending EpiSELEX-seq to measure the impact of 5hmC and 6mA on CEBP*γ* binding. (**a**) Schematic table describing the factors, library and recognition model used in analyzing the extended EpiSELEX-seq assay (c.f. Figure S3b). (**b**) K-mer enrichment analysis comparing normal and modified EpiSELEX-seq libraries, computed and displayed as in Figure S3c.

**Figure S5:**
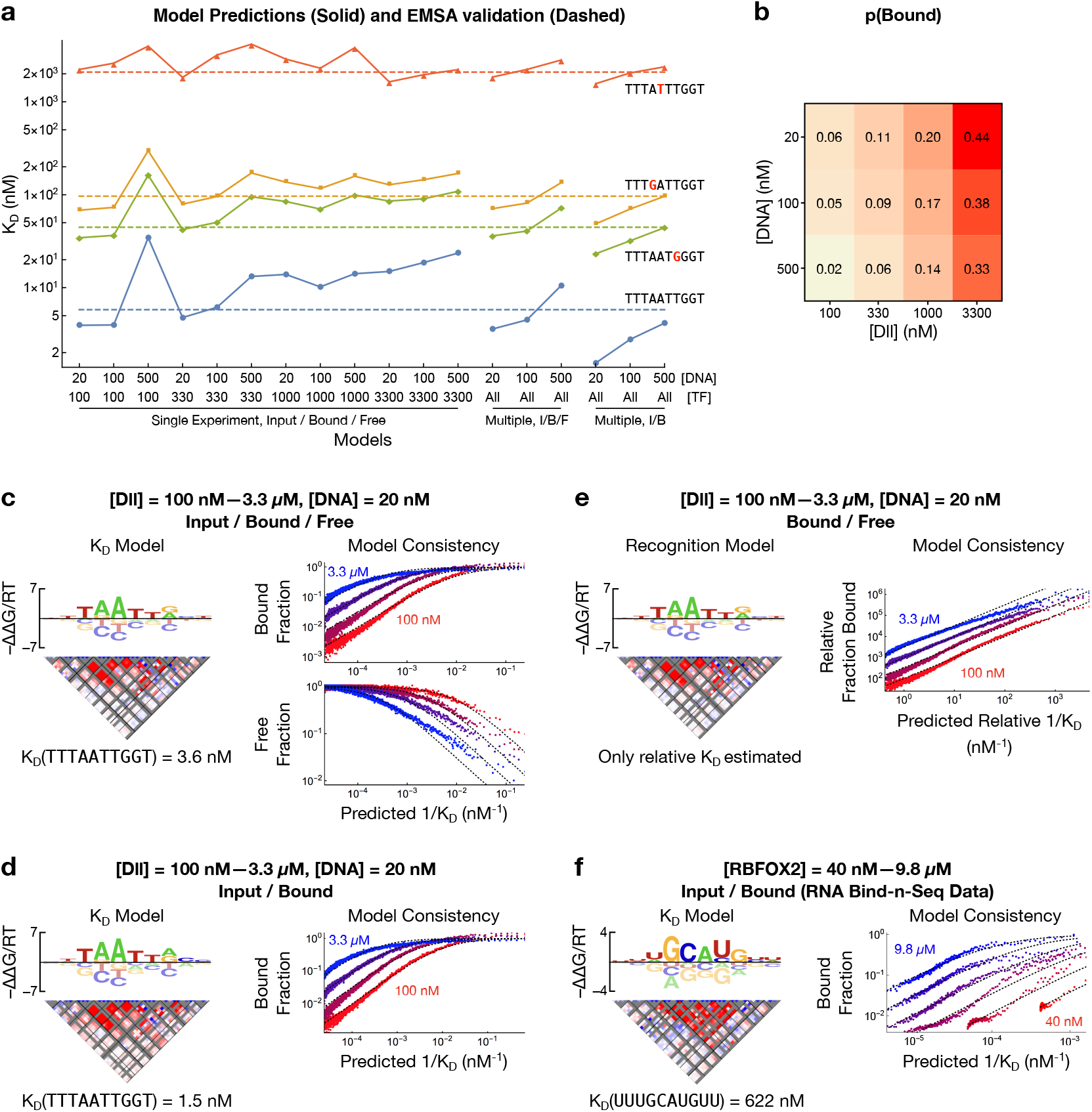
The robustness of *K_D_*-seq. **(a)** Comparison between EMSA-measured (dashed line) and different model-predicted (points) *K_D_* values for four binding probes (text). VariousInodel training strategies (x-axis) leveraged different sequencing libraries: the input/bound/free libraries fromŁ single experiment (left); the input/bound/free libraries from multiple experiments at different TF concentrations (center); or the input/bound libraries from multiple experiments at different TF concentrations (right). **(b)** Fractionlof DNA bound as inferred by ProBound when learning binding models from individual *K_D_*-seq experiments (c.f. left points in (a)). **(c)** Example *K_D_* model (left) and observed and predicted probe enrichments (right; c.f. Fig. 4c) for a model from the central points in (a). **(d)** Same as (c), but for a model from the right points in (a). **(e)** Same as (c), but only using the bound/free libraries (analogous to Spec-seq). This model can only predict relative *K_D_*, as the bound/free ratio is proportional to 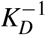 for all TF concentrations. In addition, the model predicts enrichment in the data up to a global rescaling factor. **(f)** Same as (d), but for a model derived from RNA Bind-n-Seq data for RBFOX2.

## Supplemental Methods

### Software Manual

ProBound can be run on a dedicated compute server located at probound.bussemakerlab.org. As input, this server takes a configuration file and a collection of count tables. The configurations file is in the JSON format and consists of a series of function calls:

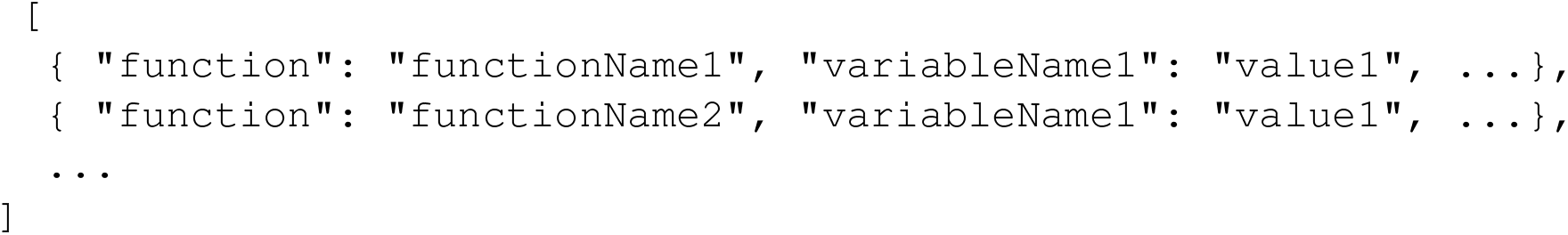

These functions configure the model components (binding modes, interactions, count tables, and enrichment models), configure the optimizer, set custom alphabets, and configure the output. For each of the components, there are functions that configure the basic parameters, functions that set custom seeding, and functions determine how the component is optimized. Below we provide documentation for these functions. In addition, we provide the configuration files that were used for the fits presented in the main text.

### Count tables

- `addTable:` This function adds a count table (containing the SELEX data) to the model.

– `countTableFile` (string, required): Path to the count table file. The table should be tab separated, have the variable region of the probe sequences in the first columns, and have the number of occurrences of each probe in each SELEX library in the following columns. This file can be gzipped. All sequences must have equal length.
– `inputFileType` (string, default is `tsv`): Format of the input file. Can either be `tsv or tsv.gz`.
– `nColumns` (integer, required): Number of columns with probe counts in the count table (that is, the first column, containing probe sequences, is not counted).
– `variableRegionLength` (integer, required): Length of the sequences in the count table.
– `rightFlank` (string, default is Specifies the constant sequence flanking the variable region to the right.
– `leftFlank` (string, default is Specifies the constant sequence flanking the variable region to the left.
– `modeledColumns` (list of integer, default is [`-1`]): Specifies what columns in the count table should be modeled. By default, all columns are included as indicated by ‘-1’.
– `transliterate` (objects, default is {“`in`”:[], “`out`”:[]}): List of edits that should be made to the probe sequences in order to encode DNA modifications. `in` lists the probe subsequence that should be substituted and `out` lists the substitutes. The lists `in` and `out` must have equal length, and each pair of sequences must have equal length.

### Enrichment models

- `addSELEX:` This function adds an enrichment model to the overall model. Enrichment models are one-to-one associated with count tables in sequential order.

– `modelType` (string, default is `SELEX`): Type of enrichment model. Possible choices are `SELEX, rhoGamma`, and `ExponentialKinetics`.
– `bindingModes` (list of integer, default is [`-1`]): Specifies what binding modes should be included in the enrichment model. [`-1`] includes all binding modes.
– `bindingModeInteractions` (list of integer, default is [`-1`]): Specifies what binding mode interactions should be included in the enrichment model. [`-1`] includes all interactions.
– `cumulativeEnrichment` (boolean, default is `true`): Specifies whether the enrichment should accumulate across columns to model repeated SELEX selection.
– `concentration` (float, default is `1`): Specifies the fixed concentration factor that multiples all activities used by the enrichment model.
– `bindingSaturation` (boolean, default is `false`): Option for `SELEX` enrichment models indicating whether the selection should be linear (∝ *x*) or or saturated (∝ *x*/(1 + *x*)).
- `enrichmentModelSeed:` This function specifies seeding of parameters for the enrichment model.

– `rho` (list of floats): Option for `rhoGamma` models seeding *ρ_r_* for each round *r*.
– `gamma` (list of floats): Option for `rhoGamma` models seeding *γ_r_* for each round *r*.
- `enrichmentModelConstraints:` Function specifying constraints on the parameters in the enrichment model and how they should be optimized.

– `fitRho` (boolean, default is `false`): Option for `rhoGamma` models specifying whether *ρ* should be optimized.
– `fitGamma` (boolean, default is `false`): Option for `rhoGamma` models specifying whether *γ* should be optimized.
– `roundSpecificRho` (boolean, default is `true`): Optionfor `rhoGamma` models specifying whether *ρ_r_* can take independent values for each round *r*.
– `roundSpecificGamma` (boolean, default is `true`): Option for `rhoGamma` models specifying whether *γ_r_* can take independent values for each round *r*.
– `trySaturation` (boolean, default is `false`): Option for `SELEX` models specifying whether the optimizer should test if setting `bindingSaturation=true` improves the model.

### Binding modes

- `addBindingMode:` This function adds a binding mode to the model and assigns it a running index, starting at 0.

– `size` (integer, required): The width of the binding mode
– `flankLength` (integer, default is `0`): Distance into the fixed flanking region that is scored by the binding mode.
– `dinucleotideDistance` (integer, default is `0`): Maximum distance between the two letters of the dimer sequence features that are included in the ΔΔ*G* model. `0` inactivates the dimer features, `1` includes only adjacent letters, such as `CG`.
– `singleStrand` (boolean, default is `false`): True indicates that only the forward should be scored and included in *Z*_bound_.
– `positionBias` (boolean, default is `false`): Indicates whether the position-bias factor should be included (*ω_a_*(*x*) is then a free) or not (*ω_a_*(*x*) = 1).
- `addNS:` Adds a non-specific binding mode (shorthand function for adding a mode with `size=0`). This function takes no arguments.
- `bindingModeSeed:` This function sets the seeding (the initial values of the parameters, before optimization) of the binding mode.

– `index` (integer): Index of the binding mode for which the seeding is specified. The seeding is applied to all binding modes if no index is specified.
– `mononucleotideIUPAC` (string): Seeds the binding mode to recognize the sequences consistent with the IUPAC string. At each position, matches get *β_a,ϕ_* = 0 and mismatches get *β_a,ϕ_* = −1.
– `mononucleotideString` (string): Seeds the binding mode to recognize sequences consistent with the string. At each position, matches get *β_a,ϕ_* = 1 and mismatches give *β_a,ϕ_* = 0. The period character (.) is a wildcard and matches any letter.
- `bindingModeConstraints:` This function specifies both constraints imposed on the binding mode during optimization and strategies used to optimize the it.

– `index, integer, required:` Index of the binding mode that will be manipulated.
– `symmetryString` (string, default is `null`): This string defines a symmetry on the binding mode. Two formats are possible:

* The first format specifies a symmetry by using letters and digits to identify equivalent positions in the binding mode. Upper and lower case letters are related through complement and digits are self-complementary. For example, the string `ab1BA` specifies a reverse-complement symmetric binding mode of size five. Here complementarity relates `a ↔ A, b ↔ B`, and `1 ↔ 1`. The string `ab1BAab1BA` specifies a 10bp binding site with a tetrameric symmetry. The pipe sign (`|`) is a barrier for dinucleotide interactions. This divides the binding mode into regions and removes dinucleotide interactions that connect different regions.
* The second format specifies a sequence of blocks that together fill in the binding mode. Each block is assigned an ID number and two block with the same ID have identical sequence recognition. A block with a negative ID is the reverse complement of a blocks with same but positive ID. Each block can be constrained to be reverse complement symmetric. For example, the symmetry string: `1:6:True` corresponds to a 6bp reverse-complement symmetric block, `1:3:False, 1:3:False` corresponds to two concatenated 3bp blocks in head-to-tail configuration, `1:3:False, −1:3:False` corresponds to a two 3bp blocks in the head-to-head configuration. Recognition of dimer sequence features that span blocks are prohibited. Note that the footprint of a binding mode cannot be modified if a symmetry is specified since the expanded binding mode would no longer have the size specified by the symmetry string.
– `roundSpecificActivity` (boolean, default is `true`): Indicates whether the binding mode activities can take different values in different SELEX rounds (columns in the count table).
– `experimentSpecificActivity` (boolean, default is `true`): Indicates whether the binding mode activities can take different values in different experiments (count tables).
– `experimentSpecificPositionBias` (boolean, default is `true`): Indicates whether the position bias parameters can take different values in different experiments. This must be `true` if the experiments have different probe lengths.
– `optimizeSize` (boolean, default is `false`): Indicates whether the size of the binding mode should be optimized. If true, the binding mode is (separately) expanded to the left and to the right, the model parameters are re-optimized, and the expanded binding mode is kept if the likelihood improved.
– `optimizeSizeHeuristic` (boolean, default is `false`): Same as `optimizeSize` but the binding mode is expanded both to the left and right (simultaneously) and the flank length is incremented.
– `optimizeFlankLength` (boolean, default is `false`): Indicates whether the flank length should be optimized. If `true`, the flank length is incremented, the model parameters are re-optimized, and the new model is kept if the likelihood improved.
– `optimizeMotifShift` (boolean, default is `false`): Indicates whether shifted versions of the binding mode should be explored. If `true`, the motif is binding model is shifted to the left and right (separately), the model parameters are re-optimized, and the new model is kept if the likelihood improved.
– `optimizeMotifShiftHeuristic` (boolean, default is `false`): Same as `optimizeMotifShift` but only a single shift is tested. This shift is found by first computing the information content for each position in the binding mode, then computing the ‘center of mass’ of the information content, and finally computing the shift such that the center of mass is at the center of the binding mode.
– `maxSize` (integer, default is `-1`): Specifies an upper limit of the binding mode size. `-1` indicates no limit.
– `maxFlankLength` (integer, default is `-1`): Specifies an upper limit of to the flank length. `-1` indicates no limit.
– `informationThreshold` (float, default is `0.1`): Threshold on the information content (computed for the first two and last two bases in the binding mode) determining whether `optimizeSize` and `optimizeSizeHeuristic` should attempt to expand the binding mode to the left and right.
– `positionBiasBinWidth` (integer, default is `1`): This setting configures the set of possible binding configurations in the probe sequence to be partitioned into bins with specified width and constrains the position-bias parameters *ω_a_*(*x*) (where *x* is a configuration) to be constant in each bin, thus reducing the number of independent parameters. By default, each bin contains a single configuration and no constraint is thus imposed.
– `fittingStages` (list of JSON objects, default is []): This setting instructs the optimizer to explore variations of the binding mode using a sequence of fitting stages. Each fitting stage can use a different set of variations and is defined by a JSON object that maps the included variations to `true`. The variations are: `optimizeSize, optimizeSizeHeuristic, optimizeFlankLength, optimizeMotifShift and optimizeMotifShiftHeuristic`.
- `symmetry:` Shorthand function for specifying the symmetry of a binding mode:

– `index` (integer, required): Specifies the index of symmetric binding mode
– `symmetryString` (string): Specifies the symmetry using the same format as in `bindingModeConstraints`.

### Interactions

- `addInteraction:` Function for adding interactions between binding modes.

– `bindingModes` (list containing two integers, required): Indices of the interacting binding modes.
– `positionBias` (boolean, default is `false`): If `true`, the binding mode interaction *ω_a_*(*x, y*) have independent value for each value of the binding mode configurations *x* and *y*. If `false`, the binding mode interaction is translationally invariant and only depends on *x* – *y* (where *x* and *y* are strand-aware coordinates).
– `maxOverlap` (integer, `0`): Maximum allowed overlap of the binding modes.
– `maxSpacing` (integer, default is `-1`): Maximum allowed spacing between the binding modes. `-1` indicates no limit.
- `interactionConstraints:` This function specifies constraints imposed on the binding mode interaction during optimization.

– `index` (integer, required): Index of the constrained binding mode interaction.
– `roundSpecificActivity` (boolean, default is `true`): Indicates whether the binding mode interaction activities can take different values in different SELEX rounds (columns in the count table).
– `experimentSpecificActivity` (boolean, default is `true`): Indicates whether the binding mode interaction activities can take different values in different experiments (count tables).
– `experimentSpecificInteraction` (boolean, default is `false`): Indicates whether the binding mode interaction can take different values in different experiments. This must be `true` if `positionBias=true` and the experiments have different probe lengths.

### General settings

- `output:` Function specifying where and how the output should be written.

– `outputPath` (string, required): Path to the output directory.
– `baseName` (string, required): String specifying the beginning of output file names (shared between all output files).
– `printTrajectory` (boolean, default is `false`): Indicates whether the optimizer trajectory should be saved.
– `verbose` (boolean, default is `false`): Indicates whether the message output to `STDOUT` should be verbose.
- `optimizerSetting:` This function configures the optimizer and accepts the following variables:

– `lambdaL2` (float, default is `1-e7`): Weight *λ* of the *L*_2_ regularizer.
– `pseudocount` (float, default is `0`): Value of *k*_Dirichlet_ (determining the weight of the Dirichlet regularizer).
– `expBound` (float, default is 4 `0`): Parameter *θ*_max_ of the exponential barrier regularizer.
– `nThreads` (integer, default is `4`): Number of threads used for parallelization.
– `nRetries` (integer, default is `3`): Number of retries that are made after numerical failures before the optimizer proceeds to the next step.
– `likelihoodThreshold` (integer, default is `0`): Smallest likelihood improvement required for a variation of a model component to be accepted.
- `lbfgsSettings:` This function specifies options for the L-BFGS optimizer.

– `memory` (integer, default is `100`): Number of previous steps kept in memory
– `maxIters` (integer, default is `500`: Maximum number of iterations.
– `convergence` (float, default is `1e-7`): Convergence criteria.
- `setAlphabet:` Function specifying the alphabet.

– `letterOrder` (string, default is `ACGT`): String specifying the set of valid letters and their order.
– `letterComplement` (string, default is `“C-G,A-T”`): String specifying what letters are mapped to each other by the complementarity transformation. The two letters in a pair are connected by a dash sign and pairs are separated by comma signs.

### Output

ProBound outputs the model parameters in the form of a JSON Object. This object has the keys:

- `countTable:` List of JSON Objects with the parameters for the count table models. Each object has the form:

– `h:` List containing the values of *h_r_* ≡ ln *η_r_*, where the index *r* runs over rounds.
- `enrichmentModel:` List of JSON Objects with the parameters for the enrichment models. The only enrichment model with parameters is `rhoGamma`:

– `rho:` List containing the values of *ρ_r_* where the index *r* runs over rounds.
– `gamma:` List containing the values of *γ_r_* where the index *r* runs over rounds.
- `bindingModes:` List of JSON Objects with the parameters for the binding modes. Each object has the form:

– `activity:` Two-level nested list containing the binding mode activities *α_e,r_*, where the indices *e* runs over experiments (count tables) and *r* runs over SELEX rounds (columns in the table).
– `mononucleotide:` Single-level list containing the mononucleotide binding mode coefficients in 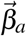 for binding mode *a*. This list can be thought of as a flattened PSAM: Letter *c* at position *x* in the PSAM has index *L* * *x* + *c*, where *L* is the length of the alphabet. For the standard alphabet this corresponds to: {*β*_*A*,1_, *β*_*C*,1_, *β*_*G*,1_, *β*_*T*,1_, *β*_*A*,2_…}.
– `dinucleotide:` Two-level list containing the dinucleotide binding mode coefficients in 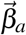 for binding mode *a*. The first index specifies the spacing between the interacting letters (`0` is `NN`, 1 is `N.N`, etc). The second index can be thought of as a flattened dinucleotide PSAM: A dinucleotide feature with letters *c*_1_ and *c*_2_ and with the first letter on position *x* has index *L*^2^*x* + *Lc*_1_ + *c*_2_, where *L* is the length of the alphabet. For the standard alphabet this corresponds to {*β*_*AA*,1_, *β*_*AC*,1_, *β*_*AG*,1_, *β*_*AT*,1_, *β*_*CA*,1_ …}.
– `positionBias:` Three-level list containing the position bias ln *ω*(*x*). The indices are: (1) experiment, (2) stand, and (3) position in the sequence. The position is specified in the 5’-3’ direction, meaning that the first position of the binding mode on the forward and reverse strands are on the opposite ends of the sequence.
- `bindingModeInteractions:` List of JSON Objects with the parameters for the binding mode interactions. Each object has the form:

– `activity:` Two-level nested list containing the binding mode interaction activities *α_e,r_*, where the indices *e* runs over experiments (count tables) and *r* runs over SELEX rounds (columns in the table).
– `positionMatrix:` Five-level list containing the binding mode interaction ln*ω*(*x,y*). The indices are: (1) experiment, (2) stand of the first binding mode, (3) strand of the second binding mode, (4) position of the first binding mode in the sequence, and (5) position of the second binding mode in the sequence. The positions are specified in the 5’-3’ direction, meaning that the first position of a binding mode on the forward and reverse strands are on the opposite ends of the sequence.

### ProBound configuration used in paper

ProBound was run with a variety of settings in order to learn the binding models shown in the figures. The corresponding JSON builder objects are provided below. These settings utilize two builder functions `addTableDB` and `oututDB`) that only work in our internal computational environment, but both these functions can be substituted. For example,

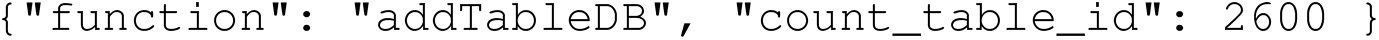

loads a count table with internal count table ID `2600` using our database. This function call should be replaced with:

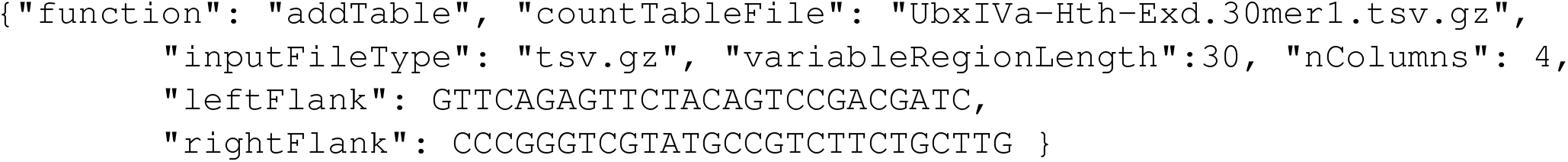

The variable values for all count tables used below can be found in Supplemental Table 2 and 3. This table also contains the accession numbers for the published sequencing data used to generate the count tables (such as `UbxIVa-Hth-Exd.30mer1.tsv.gz`). The second internal function is

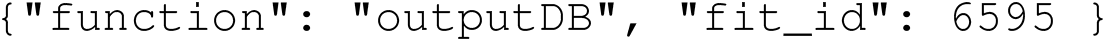

This function sets the ProBound output files using our internal database. This function call should be replaced with

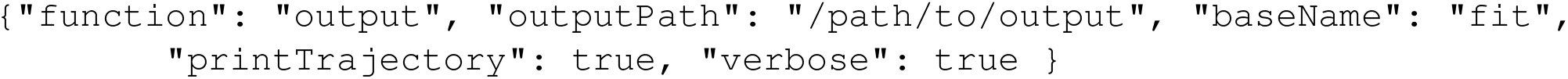

This function directs the output to the directory `“/path/to/output”` and names of the output files start with fit. Finally, some of the settings below seed the binding mode to have the sequence readout at the center. The seeding strings were based on earlier unseeded fits that are not shown. These unseeded fits explored different sizes, shifts, and flank lengths of the binding modes using `optimizeFlankLength, optimizeMotifShiftHeuristic`, and `optimizeSizeHeuristic` as illustrated by the first setting below.

### TF binding models, single-experiment

In benchmarking ProBound, each training dataset was analyzed using three settings and the best binding model was then selected based on its ability to explain the training data (see Methods). The first setting utilized one non-specific binding mode (constant across sequences) and two PSAM binding modes. The size, frame shift and flank length of the PSAM binding modes were all optimized sequentially:

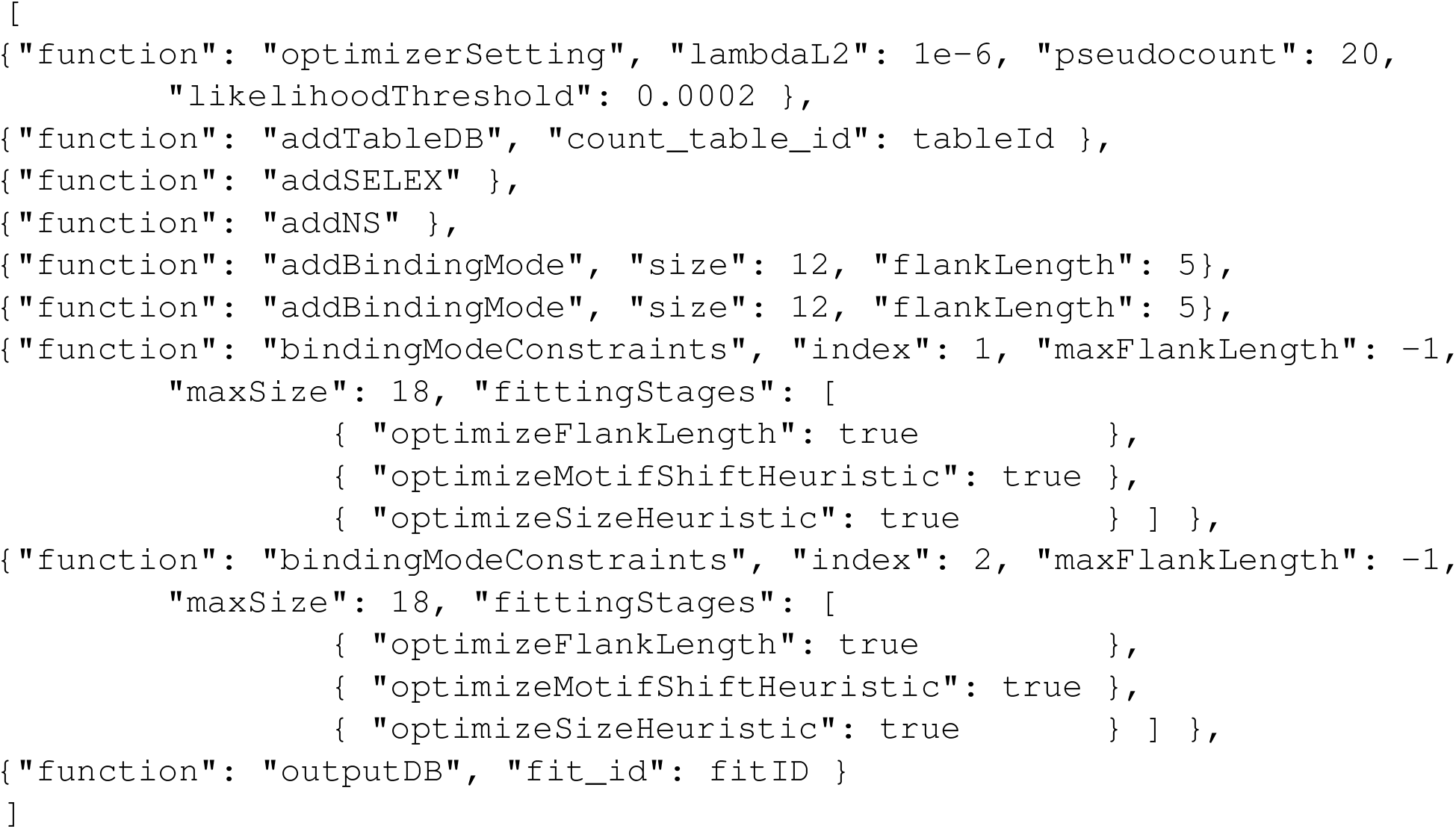

Here metadata for each count table (`variableRegionLength, nColumns, leftFlank, rightFlank`, and, when available, data accession numbers) is available in Supplemental Table 2. The second binding setting was equivalent to the first except for two changes: the non-specific binding mode was replaced by a 1bp PSAM that can absorb some sequence bias, and only the first and lasts available SELEX round was used:

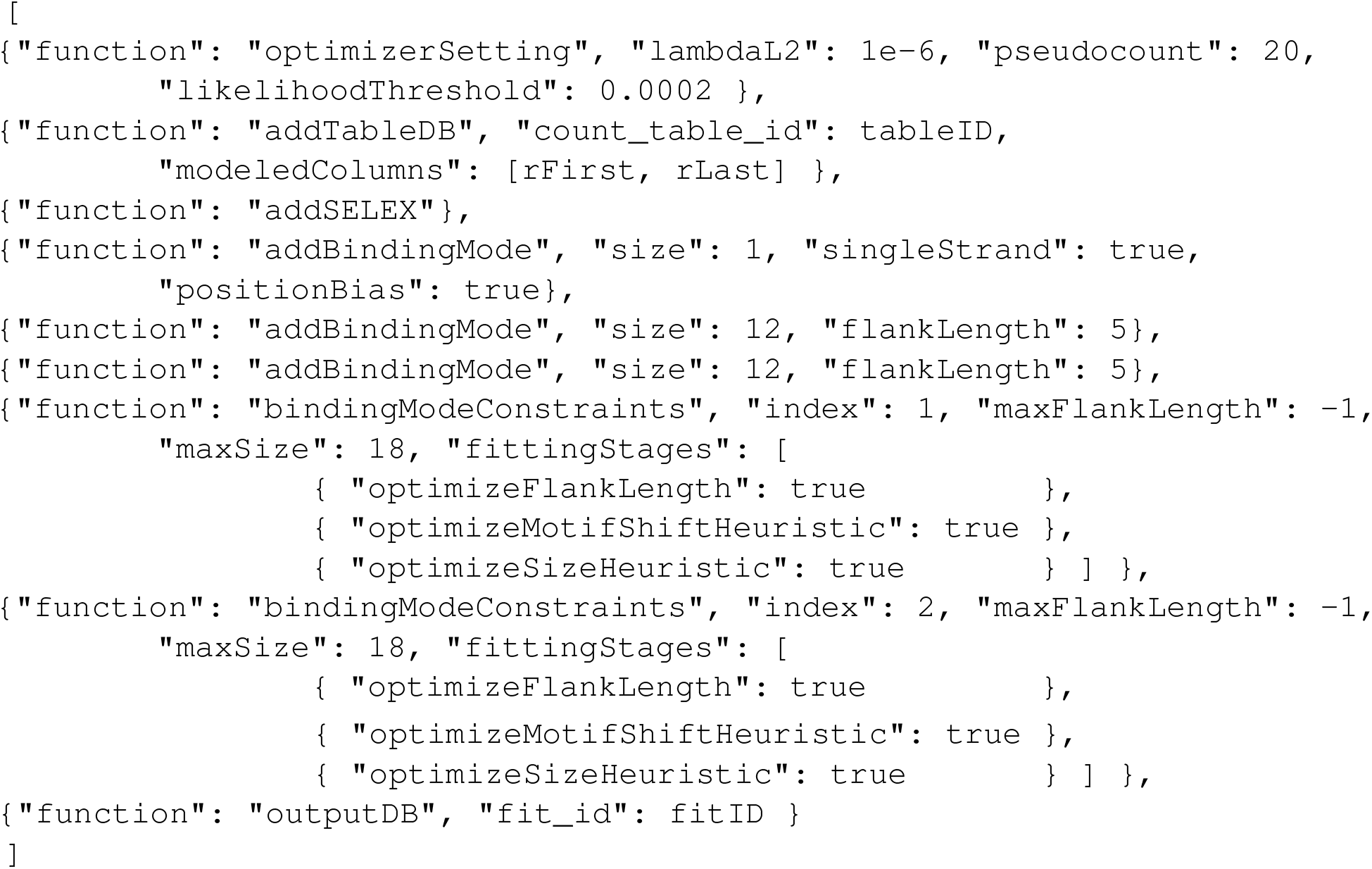

Here `rFirst` and `rLast` should be replaced with the zero-based index of the first and last available SELEX round. The third setting was also identical to the first except it learned three PSAM binding modes:

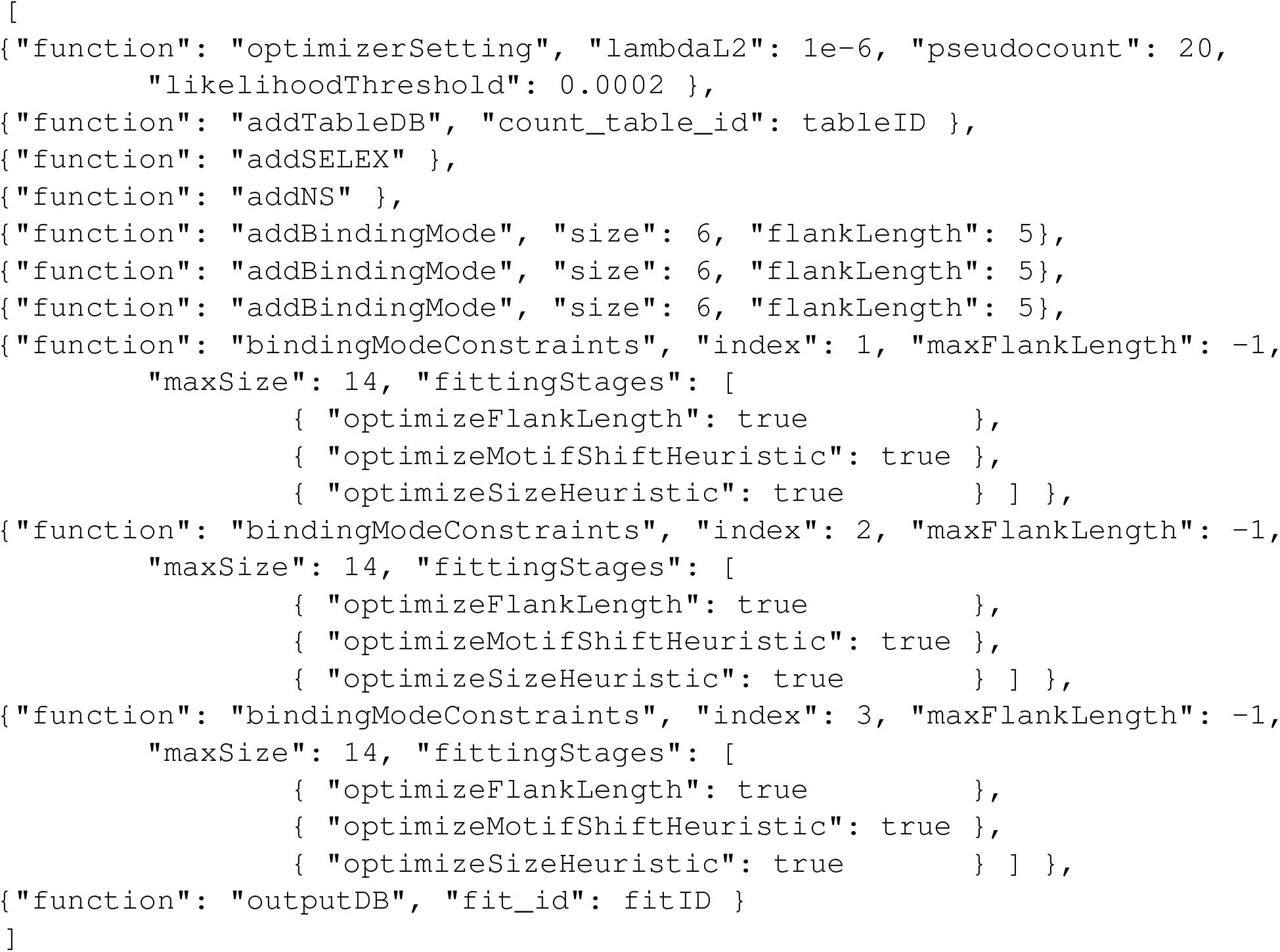

### TF binding models, multiple experiments

To learn learn a unified TF binding model from multiple SELEX datasets, the above three settings were modified to load and model multiple count tables. For example, the first setting was changed to be

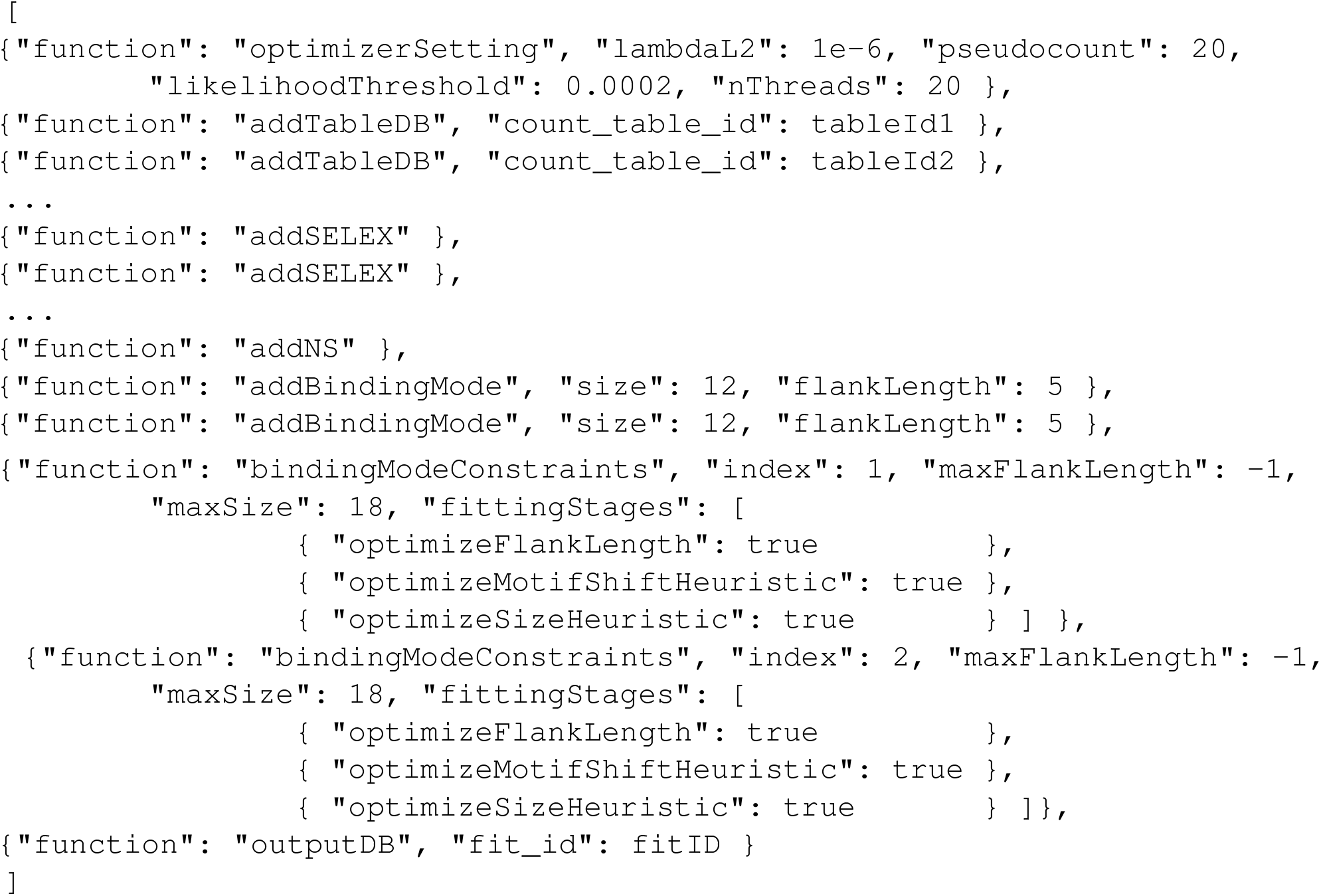

Here one call to addSELEX is added each count table loaded using `addTableDB`.

### Combinatorial SELEX

The Hth-Exd-Ubx CombSELEX-seq experiment was analyzed using following settings:

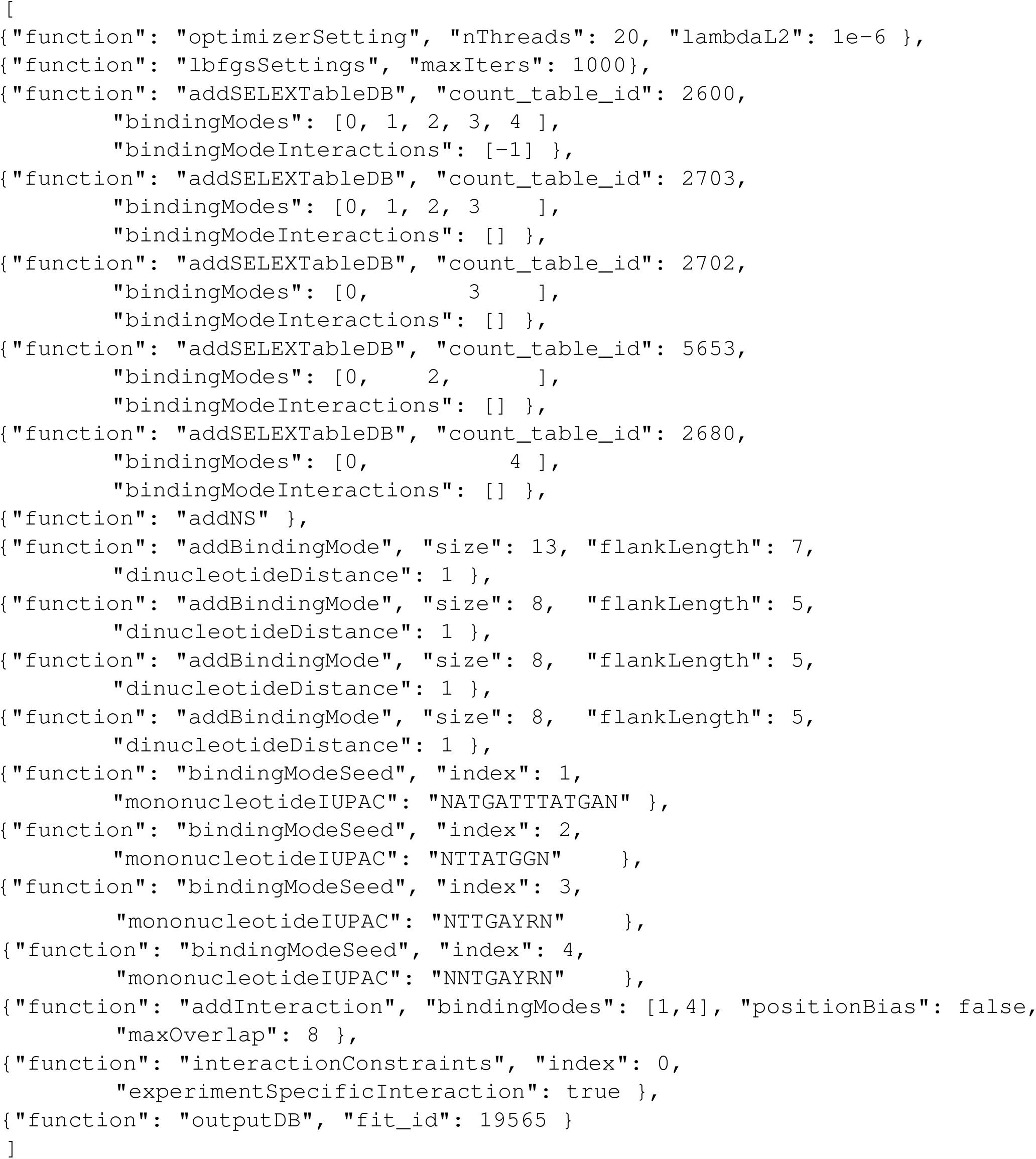

Here each SELEX enrichment model is configured to included the appropriate biding modes and interactions, as indicated in Figure S2a, The interaction corresponds to the Hth-Exd-Ubx complex. An initial unseeded fit (not shown) was used to determine consensus sequence for each TF/complex, but some modes had unfavorable offsets in the PSAMs. In the final fit (above), the PSAMs were therefore seeded to have the sequence recognition in the center.

### meCpG EpiSELEX-seq for ATF4 and CEBPγ

The meCpG EpiSELEX-seq data for ATF4/CEBP*γ* was analyzed using the following settings:

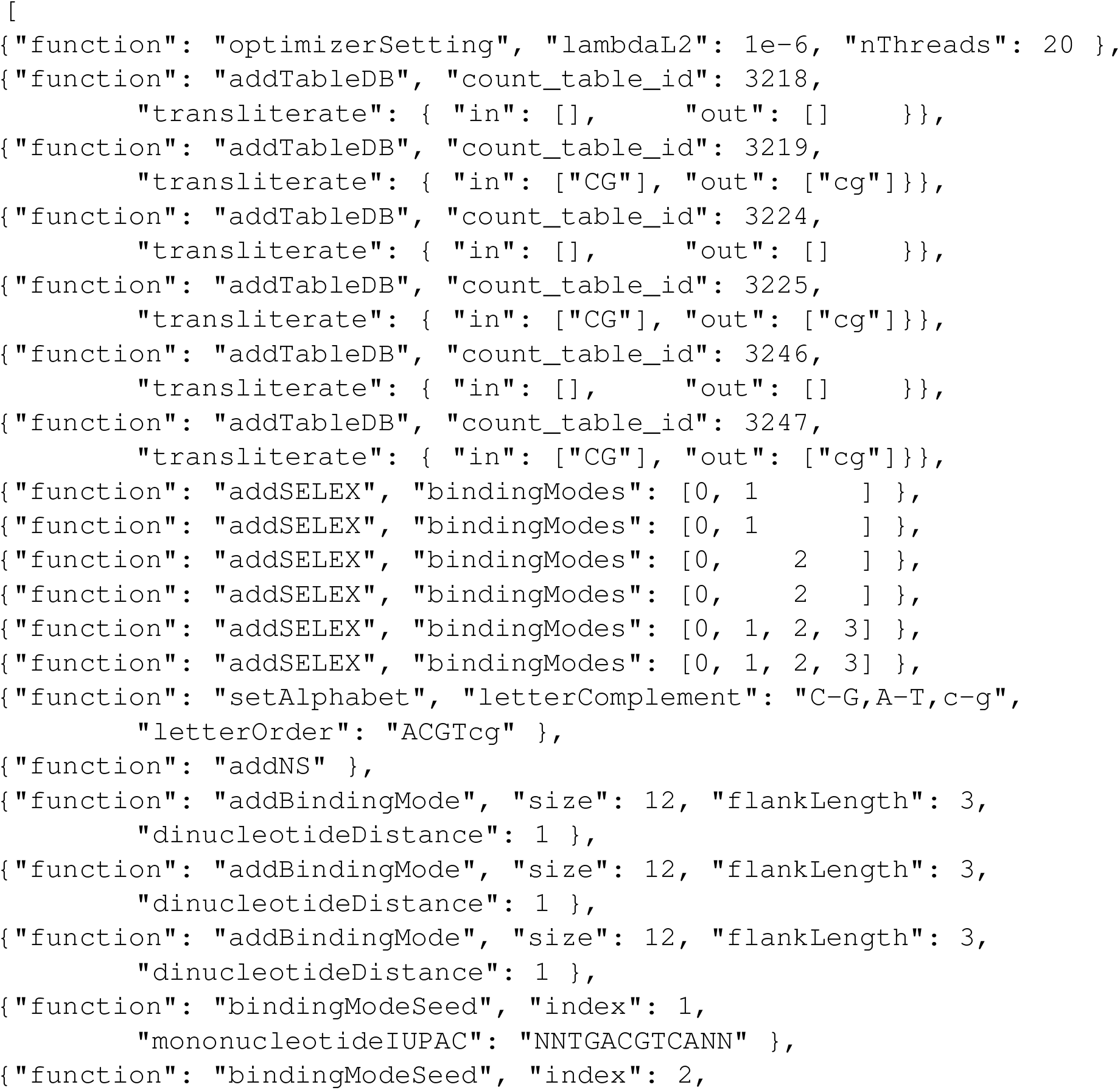
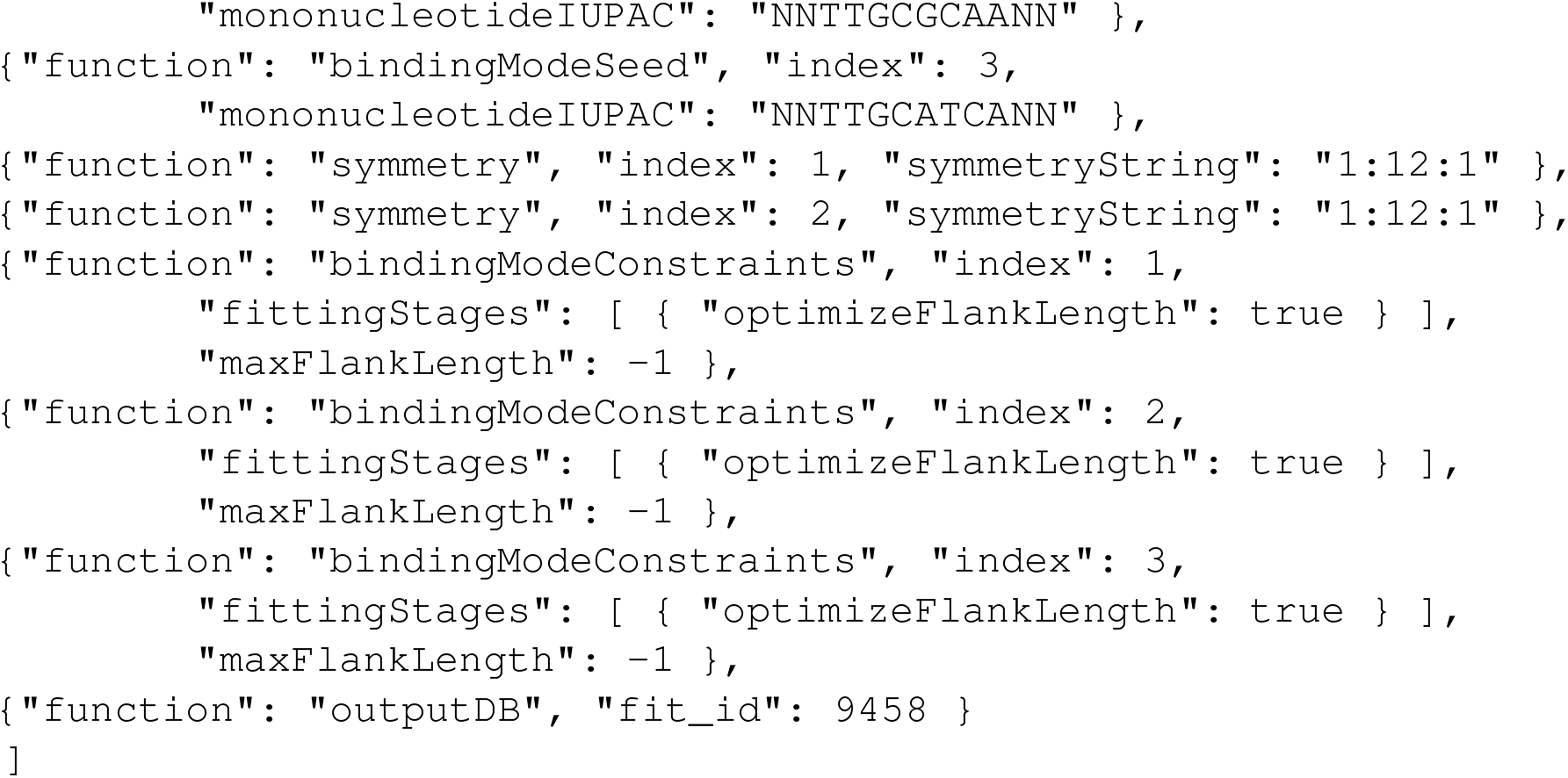

Here, only the appropriate binding modes are included in each experiment (as indicated in Figure S3b) and `CG` is transliterated to `cg` in the modified libraries to encode meCpG. The PSAMs were seeded (based on an earlier unseeded fit) to have the sequence recognition at the center, and the homodimer binding modes were constrained to be reverse-complement symmetric.

### meCpG, 5hmC and 6mA EpiSELEX-seq for CEBPγ

The meCpG-, 5hmC-, and 6mA-aware binding model for CEBP*γ* was learned using the following settings:

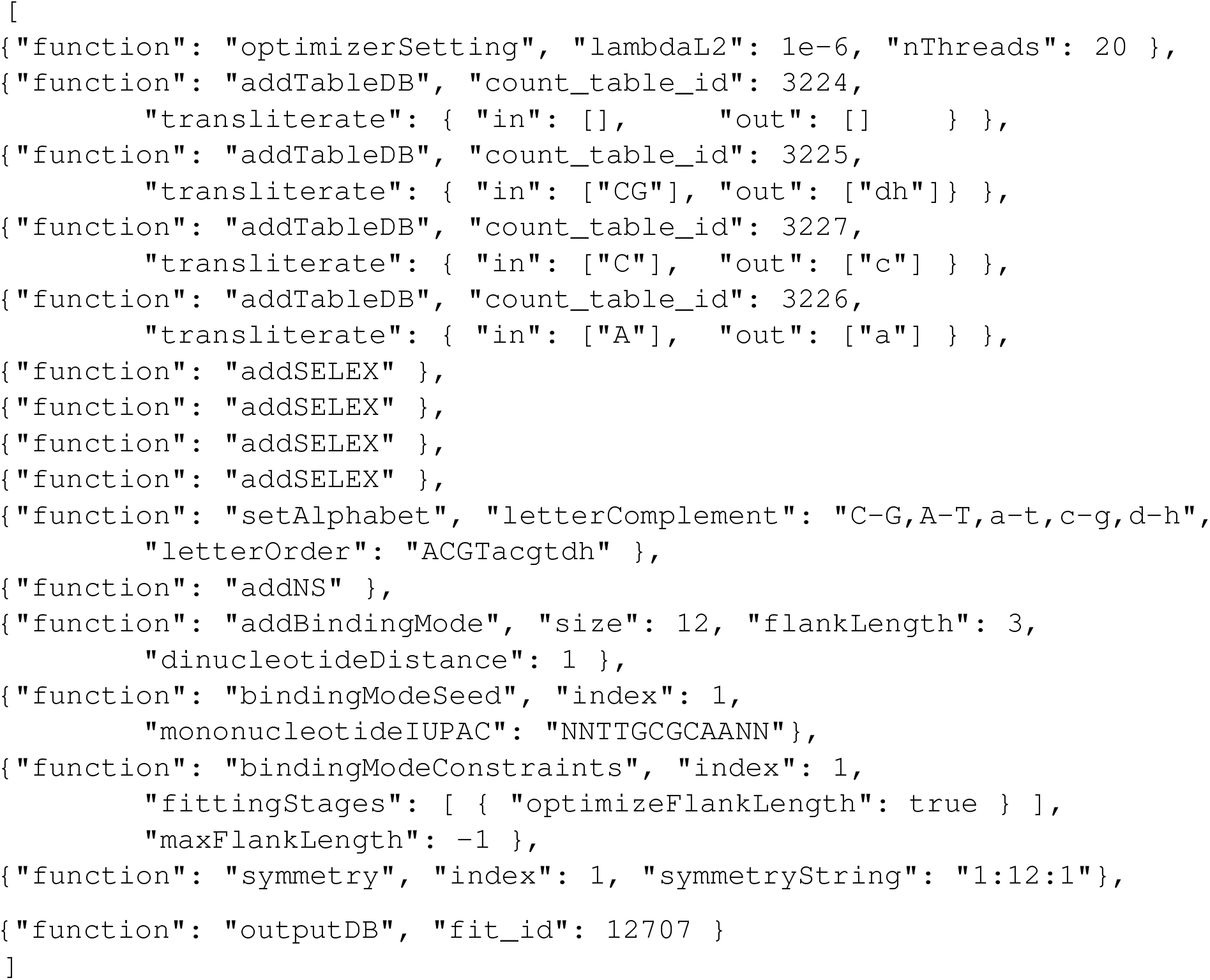

These settings encode meCpG as dh, 5hmC:G as `c` (`g` on the reverse strand), and 6mA:T as `a` (`t` on the reverse strand). While this encoding differs from that displayed in Figure S3a, it is straightforward to update the encoding of the binding model.

### RNA-binding proteins

The RNA Bind-N-seq data for RBFOX2 was analyzed using the following settings:

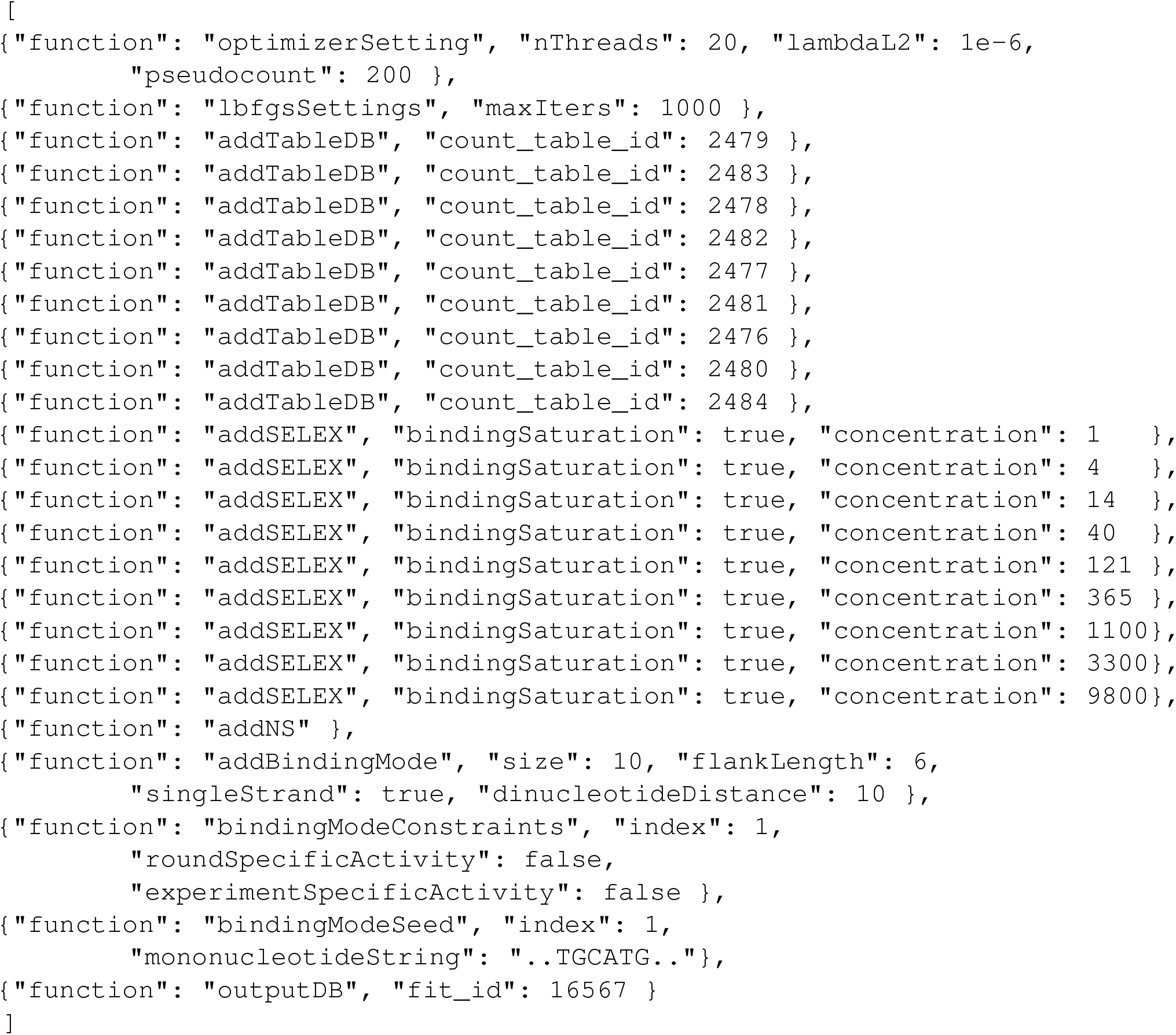

Here the SELEX model constrained the experiment-specific activities to be proportional to the RBP concentrations used in each experiment, and the binding mode was configured include all-by-all interactions and to only score the forward strand. The 1nM, 4nM and 14nM experiments have very weak binding enrichment and are not shown in Figure S5f.

### K_D_-seq - single experiment

The single-concentration *K*_D_ analyses used the following configuration:

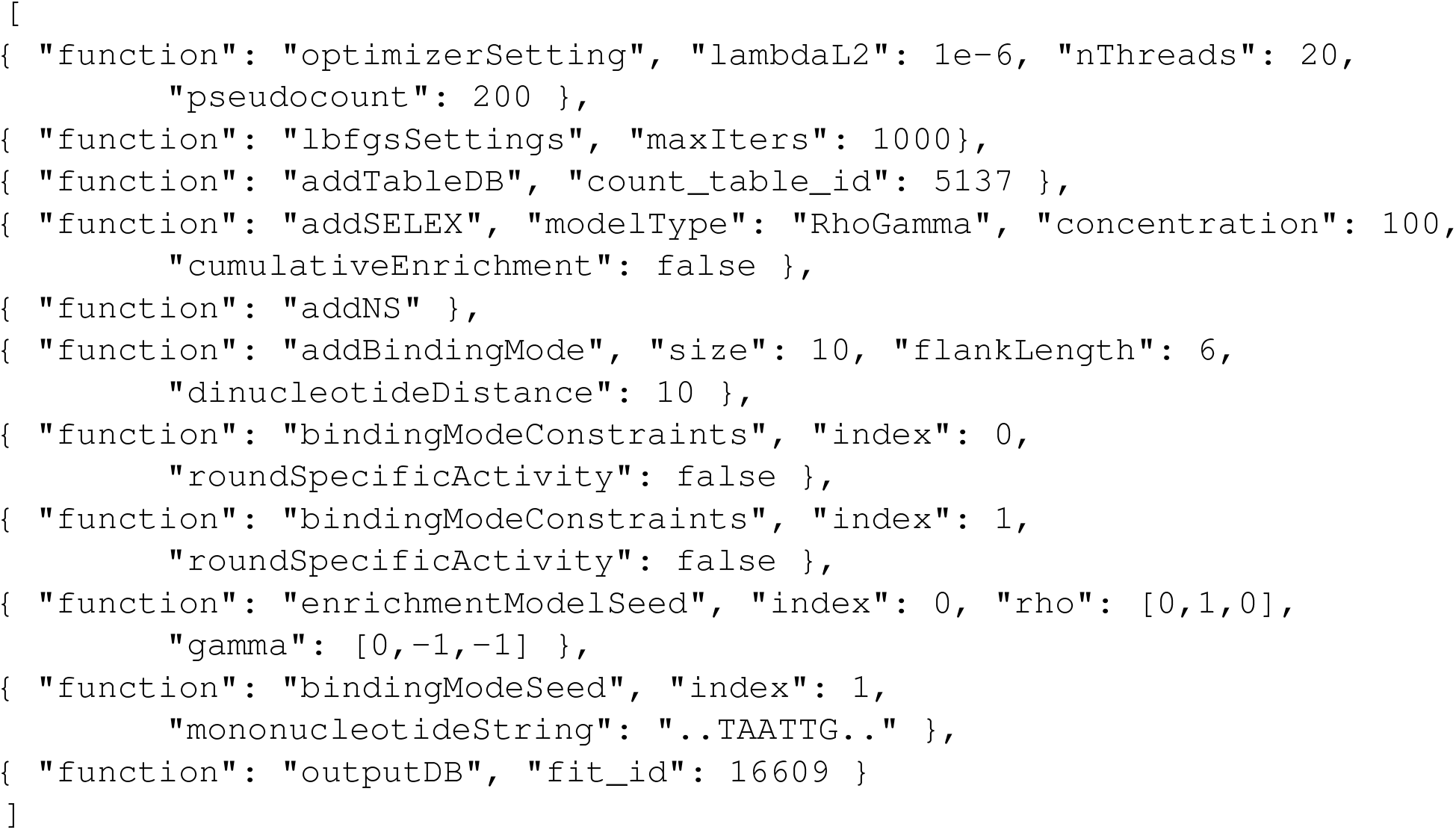

### K_D_-seq - multiple experiments

The multi-concentration KD analyses of the Input/Bound/Free libraries used the following configuration:

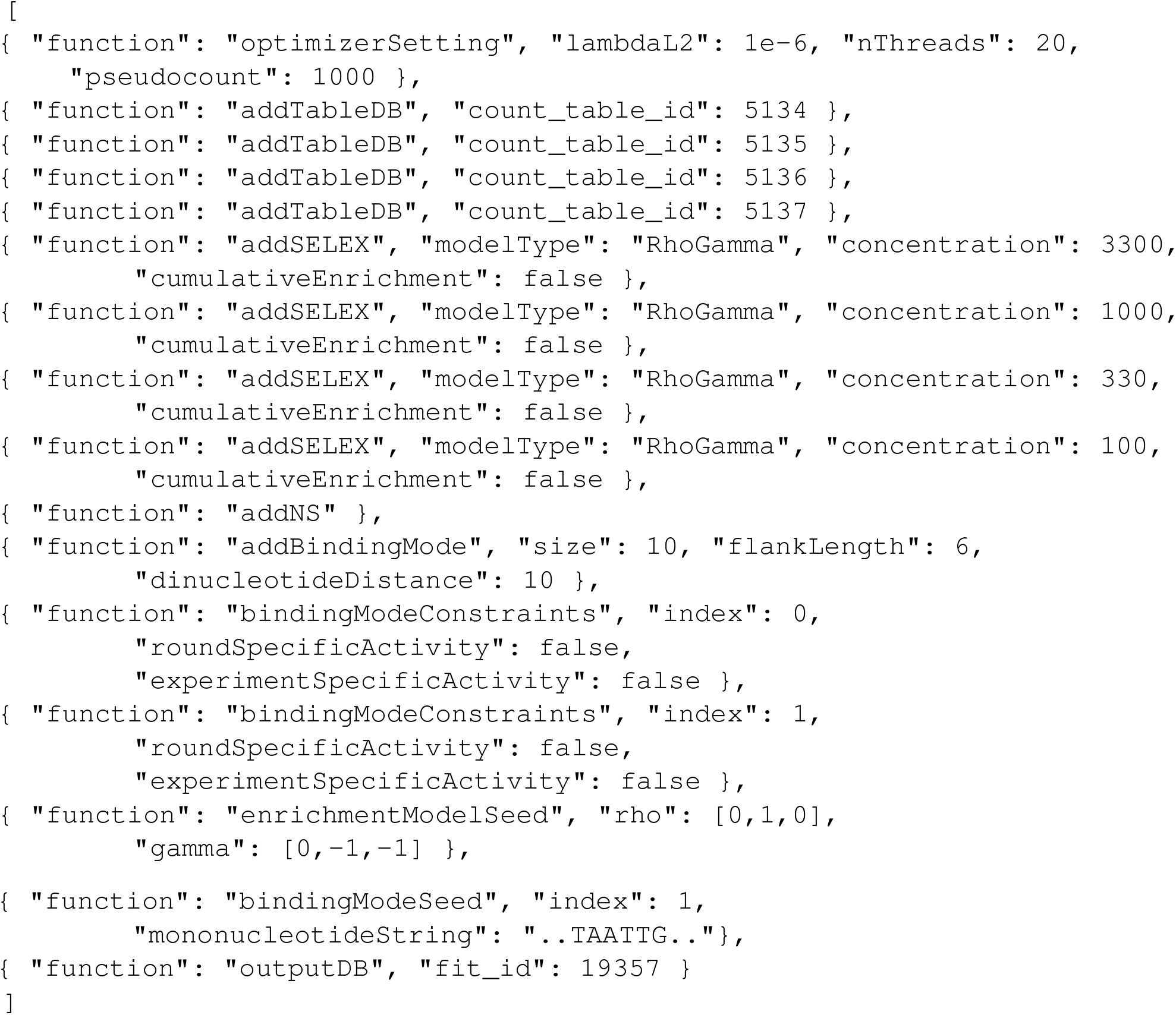

The analyses that instead analyzed the Input/Bound and Bound/Free libraries used the same configuration but with the arguments `“modeledColumns”: [0,1]` and `“modeledColumns”: [1,2]`, respectively, added to `addTableDB`.

### Peak-free ChIP-seq motif discovery - single experiment

The binding models for GR and its cofactors were learned from ChIP-seq data using the following settings:

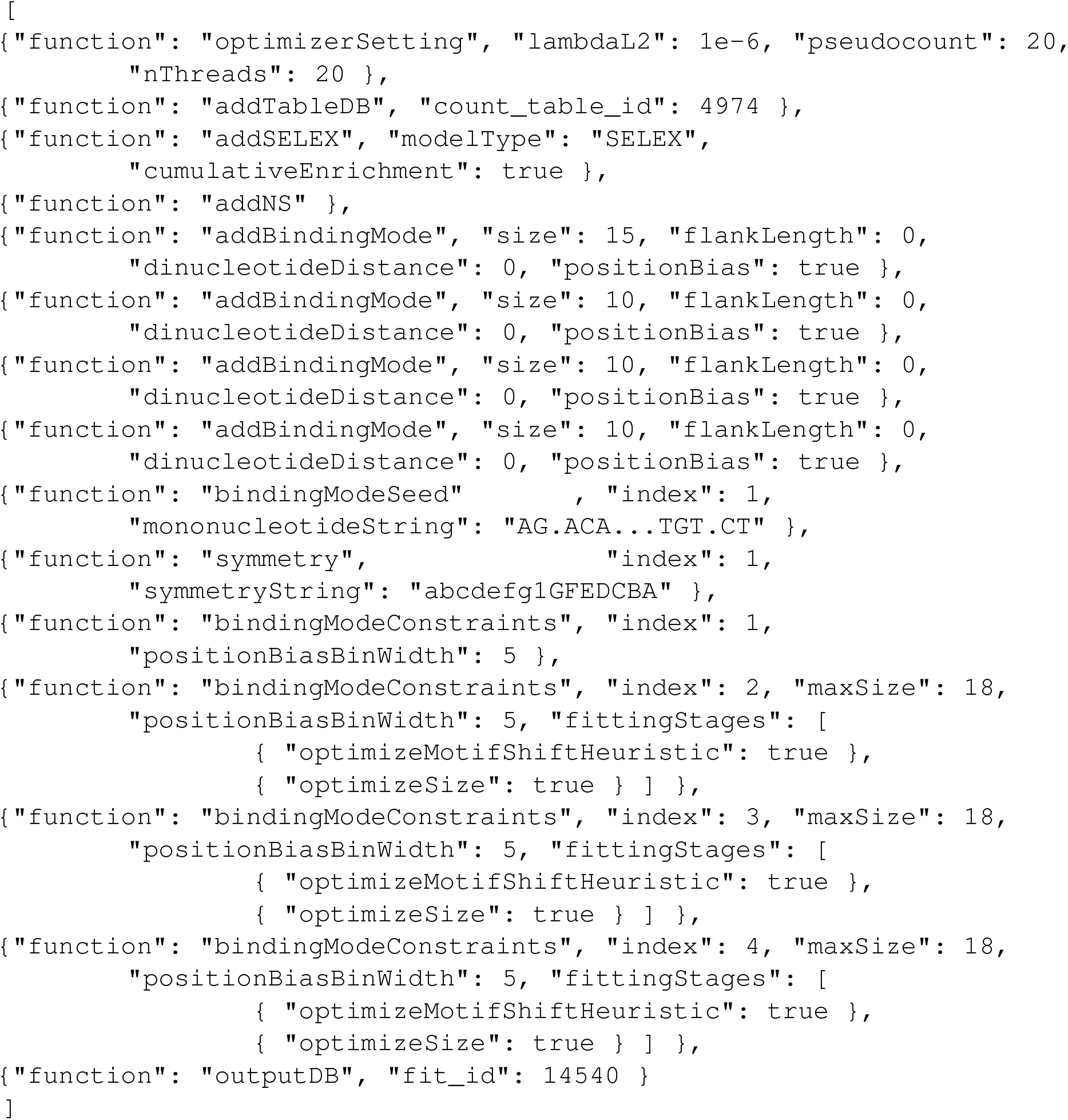

Here the GR binding mode was configured to be reverse-complement symmetric.

### Peak-free ChIP-seq motif discovery - multiple agonist treatments

The impact of CORT treatment GR binding was quantified using the following settings:

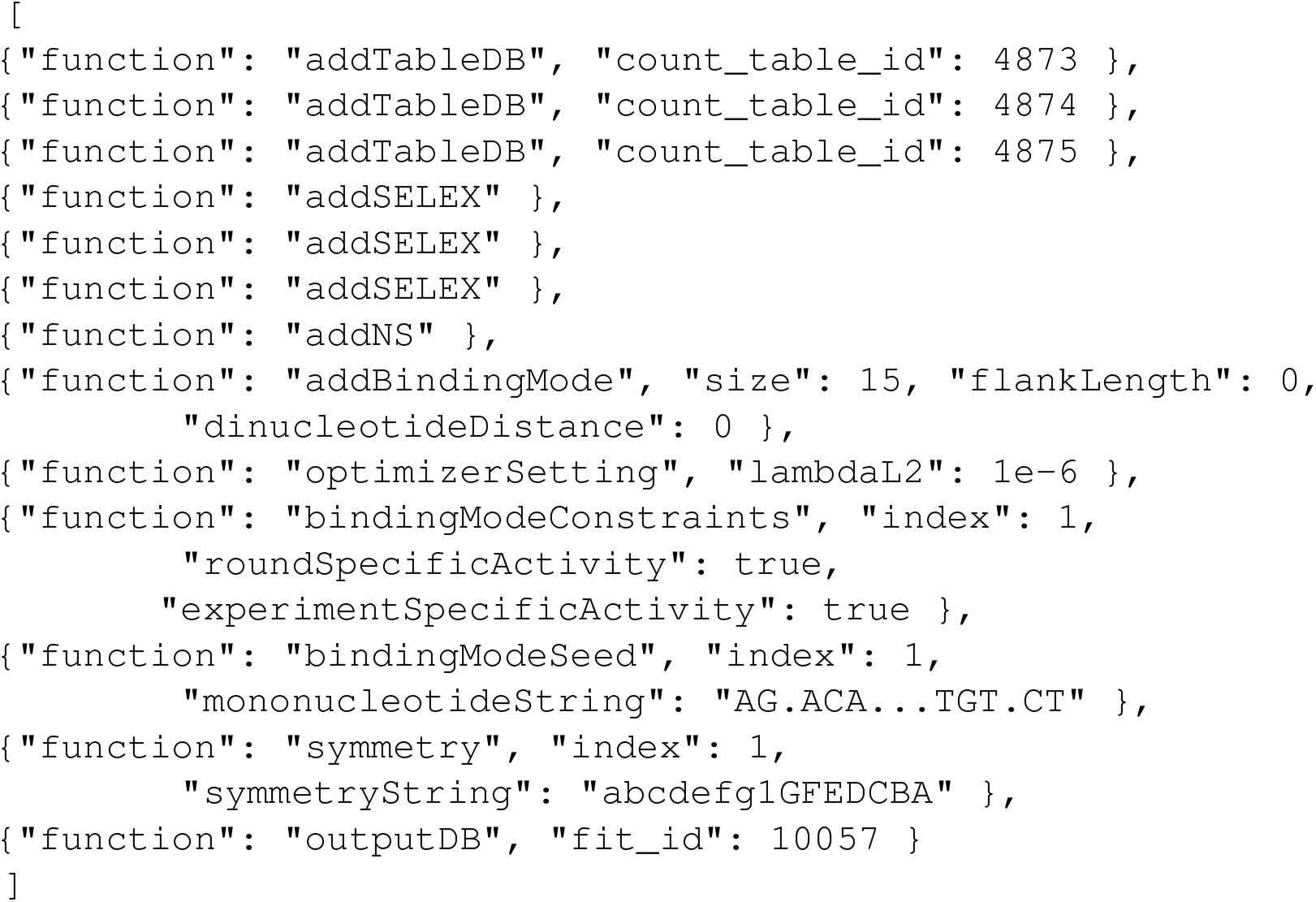

Here the binding mode is configured to have independent activities in each experiment.

### Kinase sequence specificity

The peptide-sequence specificity of tyrosine kinase Src was quantified using the following settings:

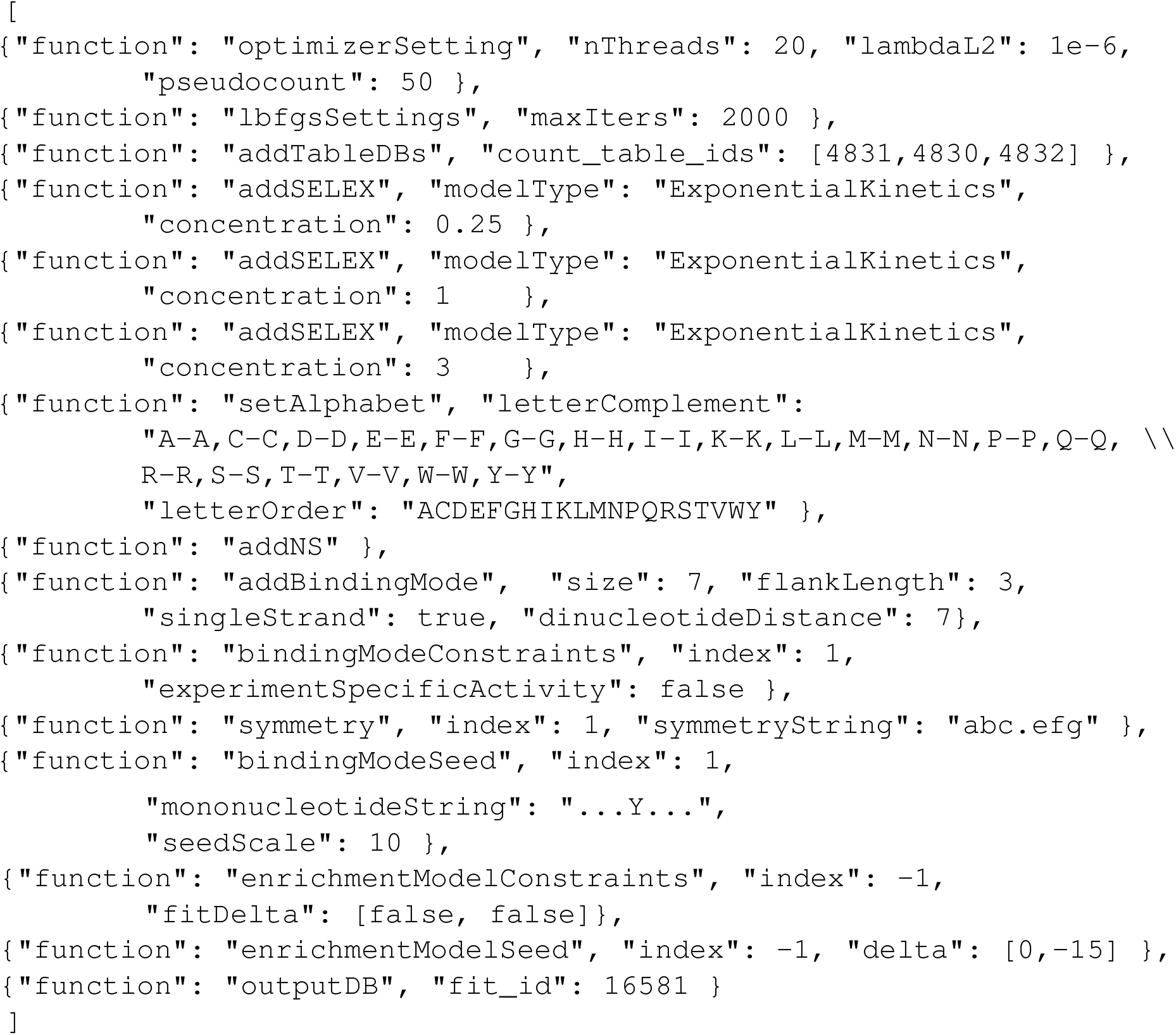

Here the `concentration` setting was used to encode the different exposures of the experiments (5min, 20min and 60min were encodes as 0.25, 1, and 3) and an extended and self-complementary alphabet was used to represent peptides. The binding mode was configured to include all-by-all interactions between the peptides and only the forward strand was scored. The commands `bindingModeSeed` and `symmetry` were used to fix the central position to recognize `Y`.

## References

1. Lambert, S. A. et al. The human transcription factors. Cell 172, 650–665 (2018).

2. Crocker, J. et al. Low affinity binding site clusters confer hox specificity and regulatory robustness. Cell 160, 191–203 (2015).

3. Farley, E. K. et al. Suboptimization of developmental enhancers. Science 350, 325–328 (2015).

4. Tanay, A. Extensive low-affinity transcriptional interactions in the yeast genome. Genome research 16, 962–972 (2006).

5. Stormo, G. D. Dna binding sites: representation and discovery. Bioinformatics 16, 16–23 (2000).

6. Zykovich, A., Korf, I. & Segal, D. J. Bind-n-seq: high-throughput analysis of in vitro protein–dna interactions using massively parallel sequencing. Nucleic acids research 37, e151–e151 (2009).

7. Zhao, Y., Granas, D. & Stormo, G. D. Inferring binding energies from selected binding sites. PLoS computational biology 5(2009).

8. Jolma, A. et al. Multiplexed massively parallel selex for characterization of human transcription factor binding specificities. Genome research 20, 861–873 (2010).

9. Isakova, A. et al. Smile-seq identifies binding motifs of single and dimeric transcription factors. Nat. methods 14, 316 (2017).

10. Slattery, M. et al. Cofactor binding evokes latent differences in dna binding specificity between hox proteins. Cell 147, 1270–1282 (2011).

11. Jolma, A. et al. Dna-dependent formation of transcription factor pairs alters their binding specificity. Nature 527, 384–388 (2015).

12. Rodriguez-Martinez, J. A., Reinke, A. W., Bhimsaria, D., Keating, A. E. & Ansari, A. Z. Combinatorial bzip dimers display complex dna-binding specificity landscapes. Elife 6, e19272 (2017).

13. Zhu, F. et al. The interaction landscape between transcription factors and the nucleosome. Nature 562, 76–81 (2018).

14. Yin, Y. et al. Impact of cytosine methylation on dna binding specificities of human transcription factors. Science 356, eaaj2239 (2017).

15. Kribelbauer, J. F. et al. Quantitative analysis of the dna methylation sensitivity of transcription factor complexes. Cell reports 19, 2383–2395 (2017).

16. Zuo, Z., Roy, B., Chang, Y. K., Granas, D. & Stormo, G. D. Measuring quantitative effects of methylation on transcription factor–dna binding affinity. Sci. advances 3, eaao1799 (2017).

17. Lambert, N. et al. Rna bind-n-seq: quantitative assessment of the sequence and structural binding specificity of rna binding proteins. Mol. cell 54, 887–900 (2014).

18. Dominguez, D. et al. Sequence, structure, and context preferences of human rna binding proteins. Mol. cell 70, 854–867 (2018).

19. Zhou, J. et al. Deep profiling of protease substrate specificity enabled by dual random and scanned human proteome substrate phage libraries. Proc. Natl. Acad. Sci. 117, 25464–25475 (2020).

20. Gee, M. H. et al. Antigen identification for orphan t cell receptors expressed on tumor-infiltrating lymphocytes. Cell 172, 549–563 (2018).

21. Ruan, S., Swamidass, S. J. & Stormo, G. D. Beesem: estimation of binding energy models using ht-selex data. Bioinformatics 33, 2288–2295 (2017).

22. Rastogi, C. et al. Accurate and sensitive quantification of protein-dna binding affinity. Proc. Natl. Acad. Sci. 115, E3692–E3701 (2018).

23. Yuan, H., Kshirsagar, M., Zamparo, L., Lu, Y. & Leslie, C. S. Bindspace decodes transcription factor binding signals by large-scale sequence embedding. Nat. methods 16, 858–861 (2019).

24. Toivonen, J. et al. Modular discovery of monomeric and dimeric transcription factor binding motifs for large data sets. Nucleic acids research 46, e44–e44 (2018).

25. Asif, M. & Orenstein, Y. Deepselex: inferring dna-binding preferences from ht-selex data using multi-class cnns. Bioinformatics 36, i634–i642 (2020).

26. Jolma, A. et al. Dna-binding specificities of human transcription factors. Cell 152, 327–339 (2013).

27. Nitta, K. R. et al. Conservation of transcription factor binding specificities across 600 million years of bilateria evolution. elife 4, e04837 (2015).

28. Yang, L. et al. Transcription factor family-specific dna shape readout revealed by quantitative specificity models. Mol. systems biology 13, 910 (2017).

29. Weirauch, M. T. et al. Evaluation of methods for modeling transcription factor sequence specificity. Nat. biotechnology 31, 126–134 (2013).

30. Alipanahi, B., Delong, A., Weirauch, M. T. & Frey, B. J. Predicting the sequence specificities of dna-and rna-binding proteins by deep learning. Nat. biotechnology 33, 831–838 (2015).

31. Davis, C. A. et al. The encyclopedia of dna elements (encode): data portal update. Nucleic acids research 46, D794–D801 (2018).

32. Khan, A. et al. Jaspar 2018: update of the open-access database of transcription factor binding profiles and its web framework. Nucleic acids research 46, D260–D266 (2018).

33. Kulakovskiy, I. V. et al. Hocomoco: towards a complete collection of transcription factor binding models for human and mouse via large-scale chip-seq analysis. Nucleic acids research 46, D252–D259 (2018).

34. Kribelbauer, J. F. et al. Context-dependent gene regulation by homeodomain transcription factor complexes revealed by shape-readout deficient proteins. Mol. Cell (2020).

35. Weber, M. et al. Distribution, silencing potential and evolutionary impact of promoter dna methylation in the human genome. Nat. genetics 39, 457–466 (2007).

36. Dantas Machado, A. C. et al. Evolving insights on how cytosine methylation affects protein-dna binding. Briefings functional genomics 14, 61–73 (2015).

37. Zhu, H., Wang, G. & Qian, J. Transcription factors as readers and effectors of dna methylation. Nat. Rev. Genet. 17, 551–565 (2016).

38. Kribelbauer, J. F., Lu, X.-J., Rohs, R., Mann, R. S. & Bussemaker, H. J. Towards a mechanistic understanding of dna methylation readout by transcription factors. J. molecular biology (2019).

39. Mann, I. K. et al. Cg methylated microarrays identify a novel methylated sequence bound by the cebpbl atf4 heterodimer that is active in vivo. Genome research 23, 988–997 (2013).

40. Kumar, S., Chinnusamy, V. & Mohapatra, T. Epigenetics of modified dna bases: 5-methylcytosine and beyond. Front. genetics 9, 640 (2018).

41. Fu, Y. et al. N6-methyldeoxyadenosine marks active transcription start sites in chlamydomonas. Cell 161, 879–892 (2015).

42. Xiao, C.-L. et al. N6-methyladenine dna modification in the human genome. Mol. cell 71, 306–318 (2018).

43. Wu, T. P. et al. Dna methylation on n 6-adenine in mammalian embryonic stem cells. Nature 532, 329–333 (2016).

44. Kriaucionis, S. & Heintz, N. The nuclear dna base 5-hydroxymethylcytosine is present in purkinje neurons and the brain. Science 324, 929–930 (2009).

45. Münzel, M. et al. Quantification of the sixth dna base hydroxymethylcytosine in the brain. Angewandte Chemie Int. Ed. 49, 5375–5377 (2010).

46. Zuo, Z. & Stormo, G. D. High-resolution specificity from dna sequencing highlights alternative modes of lac repressor binding. Genetics 198, 1329–1343 (2014).

47. Starick, S. R. et al. Chip-exo signal associated with dna-binding motifs provides insight into the genomic binding of the glucocorticoid receptor and cooperating transcription factors. Genome research 25, 825–835 (2015).

48. Polman, J. A. E., de Kloet, E. R. & Datson, N. A. Two populations of glucocorticoid receptor-binding sites in the male rat hippocampal genome. Endocrinology 154, 1832–1844 (2013).

49. Luisi, B. F. et al. Crystallographic analysis of the interaction of the glucocorticoid receptor with dna. Nature 352, 497–505 (1991).

50. Glass, C. K. Differential recognition of target genes by nuclear receptor monomers, dimers, and heterodimers. Endocr. reviews 15, 391–407 (1994).

51. Biddie, S. C. et al. Transcription factor ap1 potentiates chromatin accessibility and glucocorticoid receptor binding. Mol. cell 43, 145–155 (2011).

52. Liu, G. et al. Antibody complementarity determining region design using high-capacity machine learning. Bioinformatics 36, 2126–2133 (2020).

53. Shah, N. H., Löbel, M., Weiss, A. & Kuriyan, J. Fine-tuning of substrate preferences of the src-family kinase lck revealed through a high-throughput specificity screen. Elife 7, e35190 (2018).

54. Ryu, G.-M. et al. Genome-wide analysis to predict protein sequence variations that change phosphorylation sites or their corresponding kinases. Nucleic acids research 37, 1297–1307 (2009).

55. Hornbeck, P. V. et al. Phosphositeplus, 2014: mutations, ptms and recalibrations. Nucleic acids research 43, D512–D520 (2015).

56. Maerkl, S. J. & Quake, S. R. A systems approach to measuring the binding energy landscapes of transcription factors. Science 315, 233–237 (2007).

57. Foat, B. C., Morozov, A. V. & Bussemaker, H. J. Statistical mechanical modeling of genome-wide transcription factor occupancy data by matrixreduce. Bioinformatics 22, e141–e149 (2006).

58. Badis, G. et al. Diversity and complexity in dna recognition by transcription factors. Science 324, 1720–1723 (2009).

59. Berger, M. F. et al. Variation in homeodomain dna binding revealed by high-resolution analysis of sequence preferences. Cell 133, 1266–1276 (2008).

60. Weirauch, M. T. et al. Determination and inference of eukaryotic transcription factor sequence specificity. Cell 158, 1431–1443 (2014).

61. Zhao, Y. & Stormo, G. D. Quantitative analysis demonstrates most transcription factors require only simple models of specificity. Nat. biotechnology 29, 480–483 (2011).

62. Riley, T. R. et al. Selex-seq: a method for characterizing the complete repertoire of binding site preferences for transcription factor complexes. In Hox Genes, 255–278 (Springer, 2014).

63. Rice, J. J. & Daugherty, P. S. Directed evolution of a biterminal bacterial display scaffold enhances the display of diverse peptides. Protein Eng. Des. & Sel. 21, 435–442 (2008).

64. Shah, N. H. et al. An electrostatic selection mechanism controls sequential kinase signaling downstream of the t cell receptor. Elife 5, e20105 (2016).

65. Magoc, T. & Salzberg, S. L. Flash: fast length adjustment of short reads to improve genome assemblies. Bioinformatics 27, 2957–2963 (2011).

66. Martin, M. Cutadapt removes adapter sequences from high-throughput sequencing reads. EMBnet. journal 17, 10–12 (2011).

